# Tying down loose ends in the Chlamydomonas genome

**DOI:** 10.1101/027797

**Authors:** Frederick R. Cross

## Abstract

The *Chlamydomonas* genome has been sequenced, assembled and annotated to produce a rich resource for genetics and molecular biology in this well-studied model organism. The annotated genome is very rich in open reading frames upstream of the annotated coding sequence (‘uORFs’): almost three quarters of the assigned transcripts have at least one uORF, and frequently more than one. This is problematic with respect to the standard ‘scanning’ model for eukaryotic translation initiation. These uORFs can be grouped into three classes: class 1, initiating in-frame with the coding sequence (cds) (thus providing a potential in-frame N-terminal extension); class 2, initiating in the 5UT and terminating out-of-frame in the cds; and class 3, initiating and terminating within the 5UT. Multiple bioinformatics criteria (including analysis of Kozak consensus sequence agreement and BLASTP comparisons to the closely related *Volvox* genome, and statistical comparison to cds and to random-sequence controls) indicate that of ~4000 class 1 uORFs, approximately half are likely *in vivo* translation initiation sites. The proposed resulting N-terminal extensions in many cases will sharply alter the predicted biochemical properties of the encoded proteins. These results suggest significant modifications in ~2000 of the ~20,000 transcript models with respect to translation initiation and encoded peptides. In contrast, class 2 uORFs may be subject to purifying selection, and the existent ones (surviving selection) are likely inefficiently translated. Class 3 uORFs are remarkably similar to random sequence expectations with respect to size, number and composition and therefore may be largely selectively neutral; their very high abundance (found in more than half of transcripts, frequently with multiple uORFs per transcript) nevertheless suggests the possibility of translational regulation on a wide scale.

## Introduction

### The Chlamydomonas reference genome and annotated transcript models

The assembled Chlamydomonas reference genome is 120 Mb long, 65% GC and very repeat-rich (Merchant et al; Blaby et al). The assembly contains 17 chromosomes (~1-10 Mb) and a further 37 repeat-rich ‘scaffolds’ (0.1 -0.8 Mb). The genome has been annotated with 19,228 transcript models including transcription starts and stops, intron/exon boundaries, and coding sequence (Blaby et al. 2014), and the resulting annotated assembly is available on a public-access website (http://phytozome.jgi.doe.gov/pz/portal.html) maintained by JGI (hence ‘Phytozome’). For a subset of genes, many of these features have been verified by comparison to EST databases, as indicated on the Phytozome website. There still appears to be a need for bioinformatic methods to ‘proofread’ proposed selection of translation initiation codons in the transcript models.

### Materials and Methods

*Chlamydomonas* sequence files were downloaded from the Phytozome website, as was the .gff3 annotation file that specifies location and strand of transcript exons and coding sequence. *Volvox* predicted proteome sequences were also from the Phytozome website. BLASTP (Altschul et al., 1990) was by a local installation of the NCBI BLAST suite. Other calculations were coded in MATLAB.

## Results

### Translation start sites and 5-untranslated open reading frames

Annotating gene content from assembled genomic sequence poses many challenges (Blaby et al., 2014). There are at best weak consensus sequences for transcriptional initiation and termination/polyadenylation, so the beginnings and ends of transcripts are uncertain. RNA splicing has moderately high information-content consensus sequences, but these sequences clearly do not account for all splicing ‘choices’, and splicing intrinsically adds a huge number of degrees of freedom for computationally assembling translational open reading frames and associated 5’ and 3’ untranslated regions of transcripts. This process was aptly called ‘Gene modeling, or finding needles in a haystack’ (Blaby et al., 2014).

The genome sequence and the borders of annotated 5UT, cds, intron and 3’-untranslated regions are available on Phytozome, allowing reassembly of the complete set of 19,228 transcript models on 17 chromosomes to provide sequences of all 5UT regions and associated cds. [**Note**: in this work I chose to focus only on transcripts assigned to assembled chromosomes, leaving aside a small number of transcripts currently assigned to smaller ‘scaffolds’, since the latter are likely of less certain provenance. In addition, the 19,228 transcripts are derived from ~17,000 ‘gene’ models; the extra transcripts are due to proposed alternative initiation, splicing and or termination events. I elected to treat the transcript models as independent, since it is possible that different transcripts from some gene model might differ with respect to 5’UT or other relevant features. This provides the possibility of a minor level of duplication of results for some findings; an informal evaluation suggests that this duplication is ~randomly dispersed among functional categories].

There are no obvious bioinformatic methods to reliably determine transcriptional start sites; direct biochemical measurements (primer extension on primary transcripts and sequencing; PolII occupancy) are necessary. EST sequence comparisons provide approximate confirmation of transcription start sites in a substantial proportion of *Chlamydomonas* transcripts (Phytozome website). In the absence of other information I provisionally accept the annotated start sites as correct. These start sites, combined with annotated splicing and proposed translational start sites, result in annotated 5’ untranslated sequences (‘5UT’). For these 5UT sequences, bioinformatics approaches of various kinds can provide a quantitative appraisal of likely accuracy.

The standard model for eukaryotic translation is the ‘scanning’ model (Kozak 1978): the 40S ribosomal subunit binds at the 5’ mRNA m7GPPP cap, then scans in the 3’ direction until the first AUG, which is the translation start codon. Location of this codon triggers joining of the 60S subunit and initiation of translation (reviewed by Hinnebusch, 2011). Exceptions to this rule (skipped 5’ AUGs) may frequently be ascribed to lack of the ‘Kozak’ consensus (Kozak 1989) in inefficient initiators, which are skipped by the scanning ribosome. In some cases, a short upstream ORF (uORF) may be translated and terminated without full ribosome disengagement; provided the distance to the next AUG is not long, re-initiation can occur at a downstream AUG without rebinding to the cap. This provides the potential for regulatory mechanisms, the best-studied being yeast GCN4, where starvation effectively increases the distance the ribosome can continue scanning to reach an internal AUG (Hinnebusch 2011). Internal ribosome entry sites (IRES) are found in some viral RNAs encoding multiple polypeptides, which allow cap-independent ribosome binding and initiation; to our knowledge such sequences are rare or nonexistent outside of viral systems.

Transcripts in the annotated Chlamydomonas genome have indicated transcription start sites and 5’-untranslated regions (‘5UT’). These annotated landmarks and the reference sequence lead to the result that 13,000 of the 19228 transcript models contain one or more 5’-untranslated ATGs (Table 1).

**Table 1.** Annotated 5’-untranslated sequences for all 19228 annotated transcripts were extracted from the reference genome and analyzed for potential uORF content. For schematic of classes see Figure 1. Class 1: ATG in 5’UT sequence, in frame with reference coding sequence, without intervening stop codon; Class 2: ATG in 5’ UT sequence, out of frame with reference coding sequence, without intervening stop codon; Class 3: ATG in 5’ UT sequence, with stop codon in frame before the reference coding sequence. A given transcript can in principle have any number of each class. Sums: total transcripts containing at least one of the indicated class of uORFs.

| Number of transcript models | uORF classes present |  |  |
| --- | --- | --- | --- |
|  | 1 | 2 | 3 |
| 5628 | - | - | - |
| 6990 | - | - | + |
| 810 | - | + | - |
| 767 | - | + | + |
| 386 | + | - | - |
| 1745 | + | - | + |
| 1649 | + | + | - |
| 1253 | + | + | + |
| Sums |  |  |  |
| 5033 | + |  |  |
| 4479 |  | + |  |
| 10755 |  |  | + |

**Table 2.** Frequency of ATG in combined coding sequence (CDS), 5’UT sequence (5UT), and expected random frequency based on overall nucleotide composition of CDS and 5UT. Frequencies are numbers detected divided by total sequence length / 3. ATG_inframe: in frame with coding sequence (excluding the initiator itself); ATG_outframe: out of frame with coding sequence.

| Sequence | ATG_inframe | ATG_outframe | GC_fraction |
| --- | --- | --- | --- |
| CDS | 0.017 | 0.013 | 0.70 |
| 5UT | 0.011 | 0.021 | 0.55 |
| Rand_CDS | 0.008 | 0.016 | 0.70 |
| Rand_5UT | 0.013 | 0.027 | 0.55 |

These uORFs fall into three classes (Figure 1). Class 1 was in-frame with the annotated translation start site (henceforth, the ‘reference’ start), with no intervening stop codon. Thus, if translation initiated at the upstream ATG, an N-terminal extension (the uORF) would be appended to the expected reference peptide. Class 2 ATGs are out of frame with the reference start site, with no intervening stop codon. Initiation at class 2 ATGs thus would produce a peptide (the uORF plus frameshifted translation from the annotated coding sequence) lacking any protein sequence relationship to the predicted peptide product of the Phytozome transcript (henceforth, the ‘reference peptide’). Class 3 ATGs initiate potential 5’ uORFs that terminate *within* the annotated 5UT region. A given transcript model can have examples of all three classes of uORFs (Table 1). Note that according to the default scanning ‘1^st^ AUG’ model, class 2 and 3 uORFs should completely prevent translation of the annotated Phytozome coding sequence in ~10,000 of the 19,228 transcripts, despite the fact that in many cases this coding sequence displays high evolutionary conservation (Merchant et al. 2007); the same rule would result in an obligatory N-terminal extension to ~4000 predicted peptides encoded by class 1 transcripts. Thus these highly abundant uORFs present a *prima facie* problem with respect to translational control and the predicted proteome.

**Figure 1.**
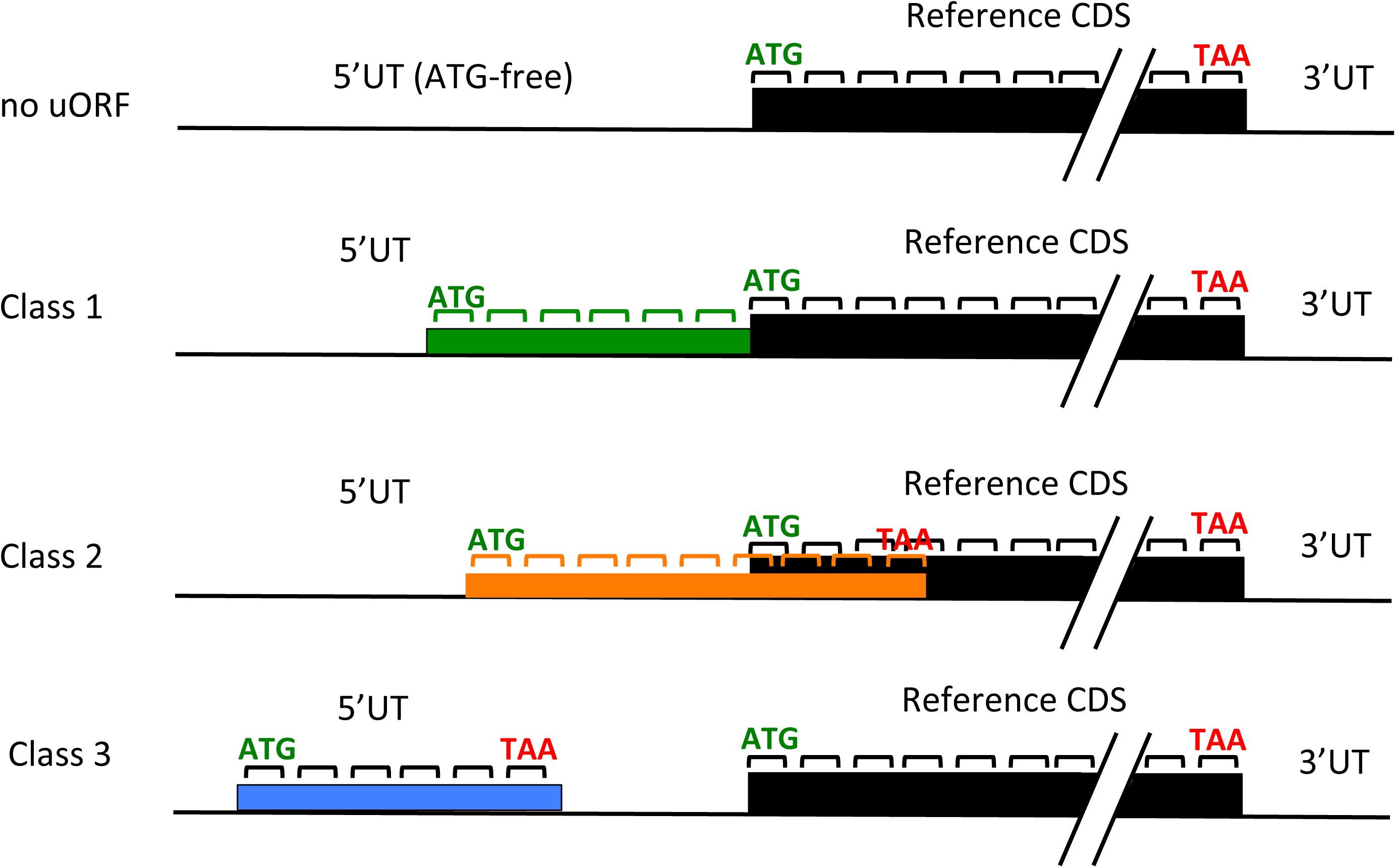
Three classes of upstream open reading frames. The 5’-untranslated sequence (5’UT)can contain no ATG in any frame (top), resulting in no upstream open reading frame (‘uORF’). It can contain one or more ATGs in frame with the main coding sequence (‘Reference CDS’), without an intervening stop codon (Class 1). Class 2 is the same as Class 1 but in a different reading frame; in general the Class 2 uORF will terminate shortly after entering the main coding sequence out-of-frame. Class 3 initiates (in any frame) and terminates within the 5’UT. Note that we consider a maximum of one Class 1 and one Class 2 uORF per transcript, although these uORFs can contain internal ATGs that could in principle initiate a ‘different’ Class 1 or Class 2 uORF. A given transcript can (and frequently does) contain multiple Class 3 uORFs, in the same or different frames. We consider all of these as separate individuals.

In a broad range of organisms, ATG frequency is significantly reduced in 5’untranslated sequences compared to coding sequences (Zur and Tuller, 2013). In the Chlamydomonas annotation, the frequency of ATGs in frame with coding sequence is one-third lower in 5’UT than in coding sequence (excluding the reference initiator itself), while the frequencies of out-of-frame ATGs is about 50% higher. These departures are largely due to deviations from random expectations (based on overall nucleotide frequencies), specifically in the coding sequence. Overall, though, ATG frequency in 5UT and coding sequences are nearly identical (0.032 vs 0.030), in contrast to results in other organisms (Zur and Tuller, 2013).

### Class 1 uORFs are longer than expected for random sequence

If uORFs have biological relevance, this should likely be reflected in statistical sequence characteristics. To make this determination a control sequence set is needed. First, a ‘scrambled 5UT-ome’ was constructed by individually randomizing sequence of each 5UT sequence, thus preserving nucleotide composition but not sequence. This control set of uORFs will reflect sequence-independent consequences of the nucleotide composition and length distribution of the reference 5UTs. A more stringent control for sequence dependence was derived by ‘mutagenizing’ the annotated 5UTs by randomly replacing on average one tenth, one fifth or one half of the nucleotides in each 5UT with another nucleotide chosen based on the overall 5UT nucleotide frequency. These controls will lack only strongly sequence-dependent features when compared to the reference 5UTs.

Results of these comparisons with respect to number and length of uORFs were very clear, and strikingly different for the three classes. For classes 2 and 3, uORF lengths were essentially identical for the reference and the controls (either random or mutated; Figure 2A). In contrast, class 1 uORFs were significantly longer in the reference than in the controls. Mutation of one in five nucleotides was almost as effective at eliminating sequence dependence as complete randomization, suggesting a high degree of sequence dependence in the real sequence. The numbers of classes 1 and 3 uORFs were approximately similar in reference and controls (Figure 2B); however, class 2 uORFs doubled in abundance in the randomized control compared to the reference.

**Figure 2.**
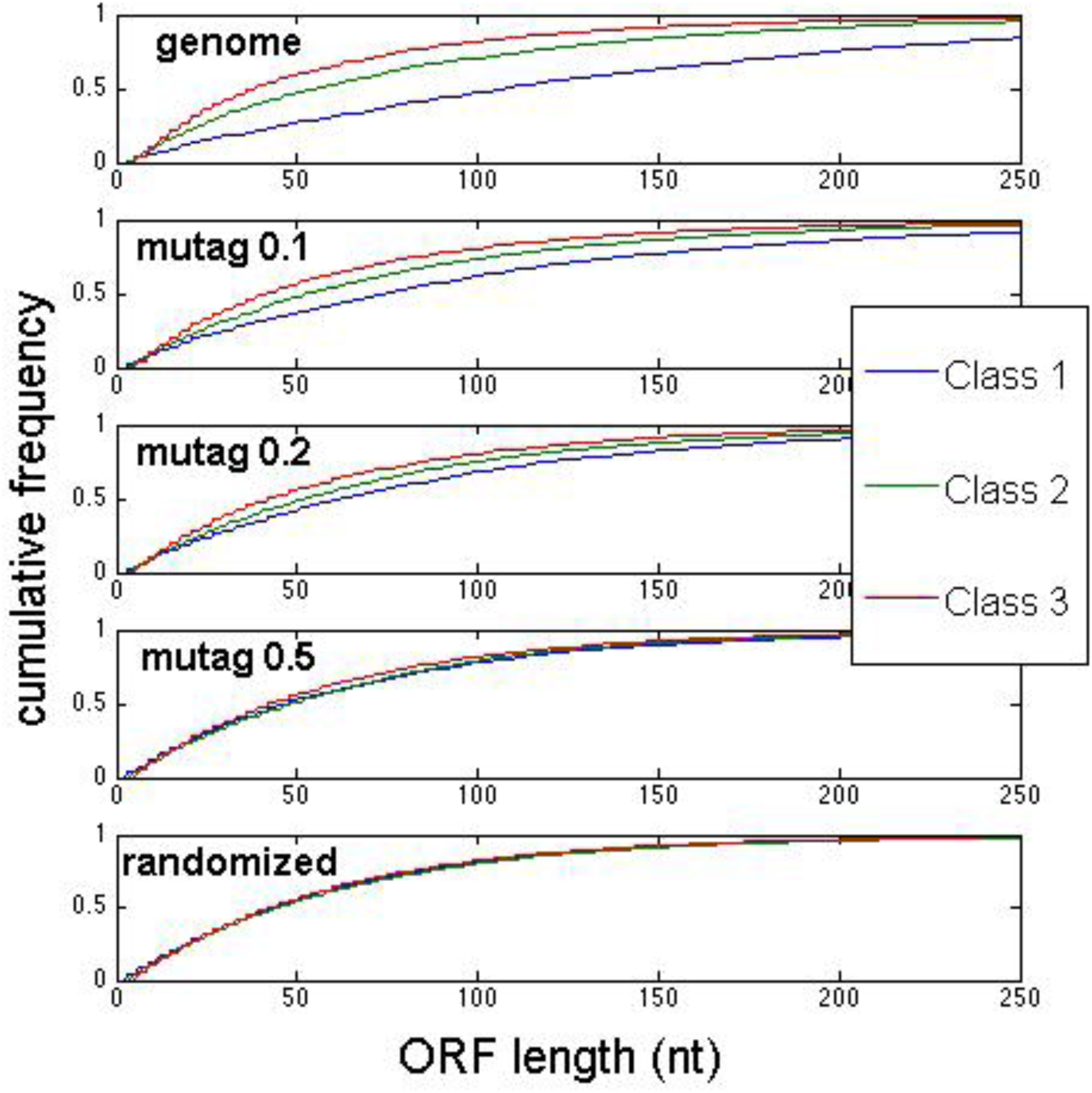
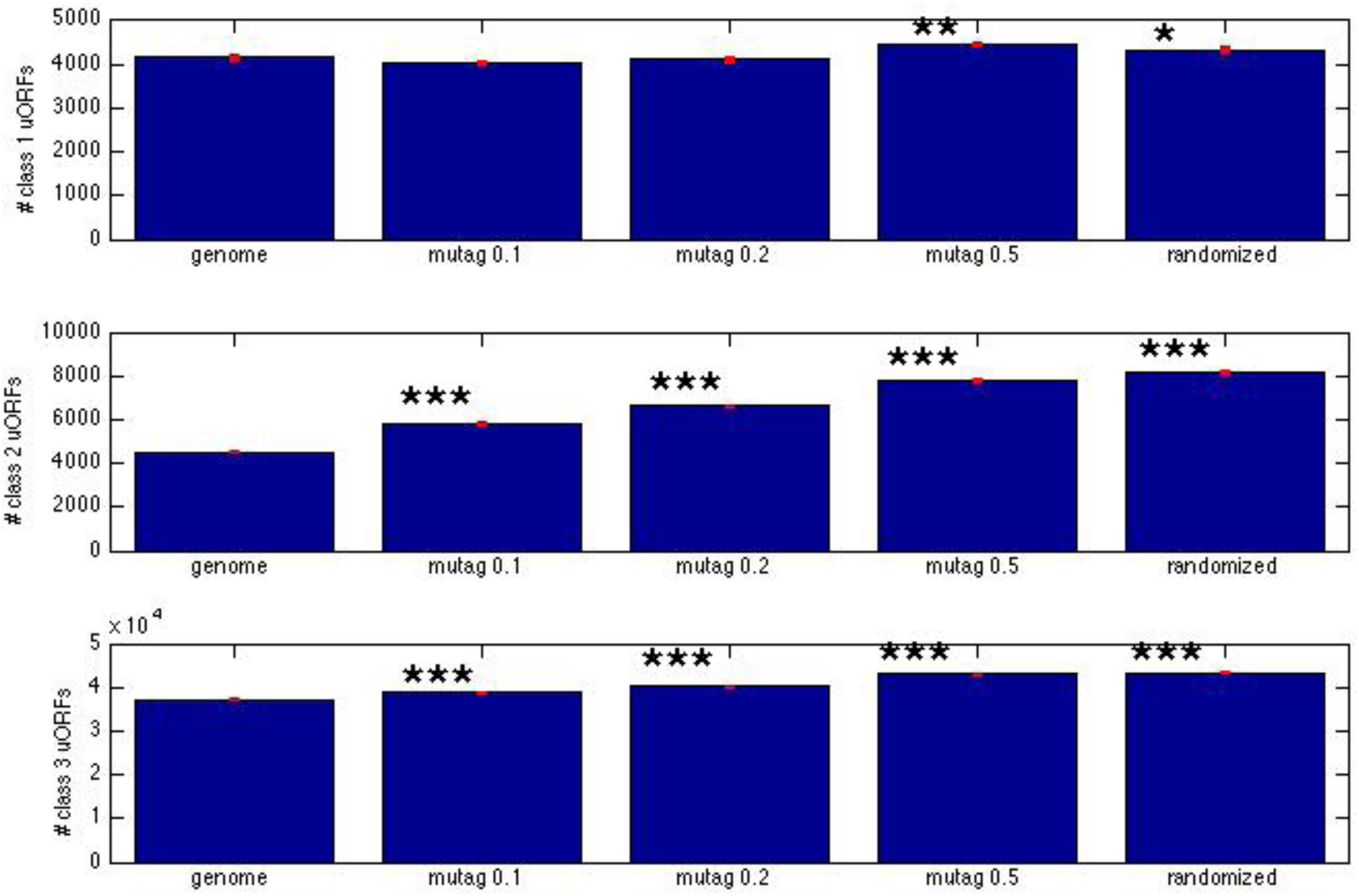
Statistics of uORF length and number in the genome, and in partially or fully randomized controls. A: Class 1 but not Class 2 or Class 3 uORFs are longer than the random expectation. The cumulative length distribution of uORFs is shown (‘genome’, top). Lower panels: the set of 5’UT sequences was partially or fully randomized (by replacing 1 in 10, 1 in 5, 1 in 2, or all nucleotides in each 5’UT with random selections from the overall nucleotide frequency distribution of the complete collection of 5’UT sequences). (Note: the randomized distribution for all classes is essentially identical to the class 3 length distribution for the actual genomic Class 3 sequences). **B: Total numbers of uORFs with and without randomization.** The small red bar represents a hypothetical standard deviation based on the assumption that numbers in each category are Poisson-distributed (square root of the number observed). Stars represent P-values for a t-test comparing each randomization to the genome, using these standard deviations: ^*^, P<0.05; ^**^: P<0.01; ^***^: P<0.001).

Note that the random expectation is that class 2 should be twice as abundant as class 1, since there are two ‘wrong’ frames and one ‘right’ frame.

These results suggest sequence-dependent constraints that (1) preserve class 1 uORFs at significantly longer than expected by chance; (2) suppress the numbers of class 2 uORFs to about half the level expected by chance. Class 3 uORF numbers and lengths are strikingly well predicted by nothing more than 5UT nucleotide composition and length distribution, and thus exhibit no sequence dependence detectable by this bulk approach. The close statistical correspondence of the genomic Class 3 uORFs and those constructed from randomized sequence suggests that most Class 3 uORFs are effectively neutral sequence (though statistically significant increases Class 3 uORF numbers in randomized controls [Figure 2B] suggests purifying selection over at least a subset). The suppression of Class 2 uORF number relative to random controls suggests that Class 2 uORFs, in contrast, are subject to significant purifying selection. This could be understood based on the scanning model for translation initiation since initiation from a Class 2 AUG would block even post-termination re-initiation at the reference AUG, since scanning is probably generally (though perhaps not exclusively) unidirectional (Hinnebusch 2011), and class 2 termination occurs 3’ to the reference initiation site (Figure 1). Thus class 2 AUG’s could be particularly damaging to expression of the main coding sequence.

### A Chlamydomonas Kozak consensus

These observations raise issues with respect to the standard translation model. Since the reference initiation ATG generally starts translation of a long peptide, which is frequently conserved across species, it is highly unlikely that class 2 and class 3 ATGs are exclusive sites of initiation. This suggests a high level of selectivity, since nearly 2/3 of transcripts are assigned class 2 and/or class 3 uORFs – so if the reference ATG is in fact the one used in vivo, multiple 5’ ATGs must be skipped, or must fail ribosome disengagement in a majority of transcripts – thus, the scanning model would be the exception rather than the rule.

There are two obvious escapes from this problem. The simplest is if the annotated transcription start site is misplaced at a position 5’ of the real start. For a substantial subset of Phytozome gene models, there is evidence from EST sequence that the transcript does indeed cover some or all of the proposed 5’UT, so this is unlikely to be the entire explanation.

Another escape would be strong sequence constraints suppressing initiation at uORFs. This is a known mechanism in other eukaryotes, where absence of a ‘Kozak’ consensus sequence can allow skipping of a 5’ AUG (Kozak 1989; Hinnebusch 2011). Using the Weblogo calculation (Crooks et al 2004) with all 19,228 reference initiator ATGs yielded a consensus with striking similarity to the Kozak consensus found for human mRNAs (Figure 3).

**Figure 3.**
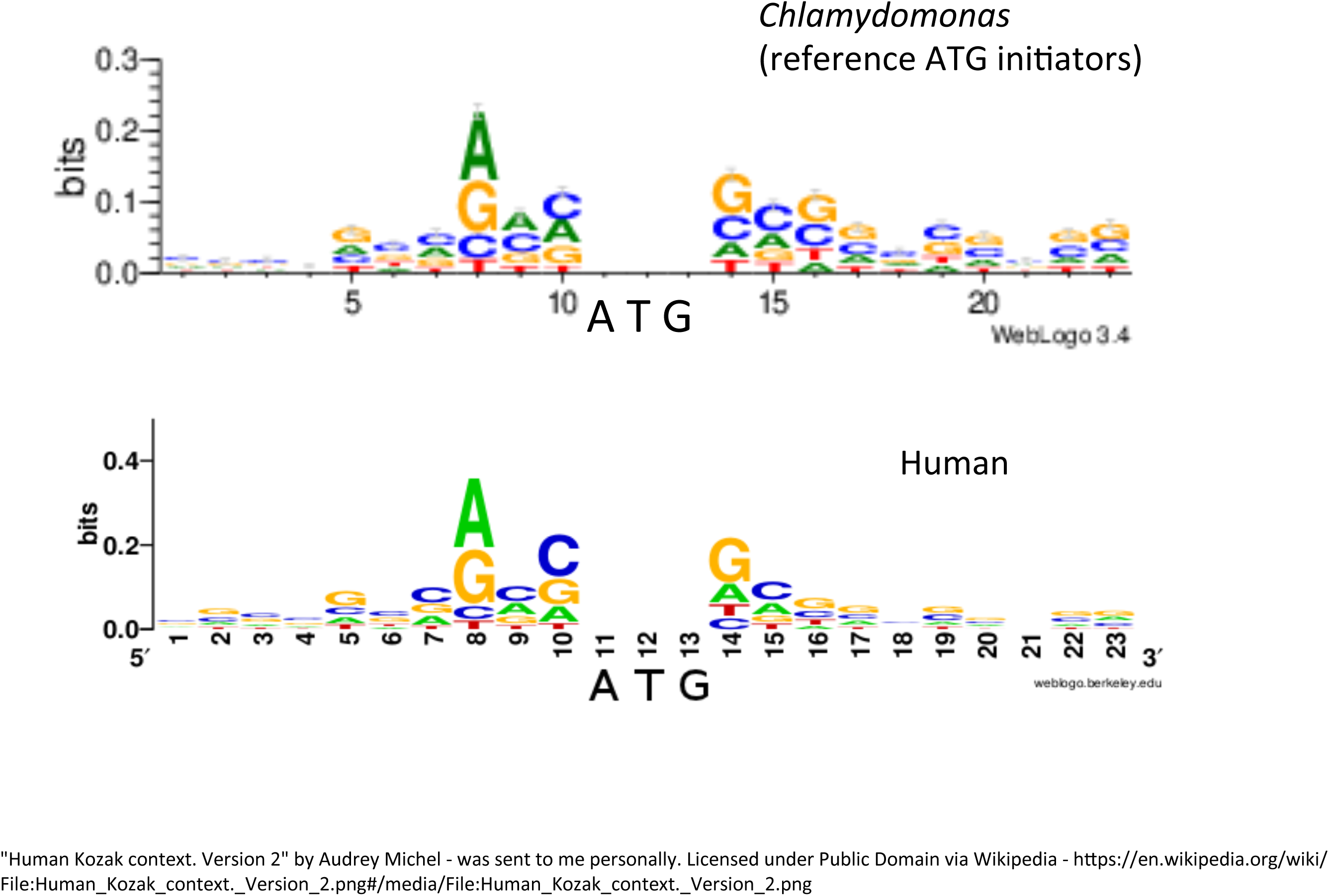
Kozak-like consensus sequence around *Chlamydomonas* reference initiator ATGs. All 19,228 sequences were fed to the online WebLogo tool (http://weblogo.threeplusone.com) (Crooks et al., 2004). A comparable plot for human mRNAs was downloaded from Wikipedia (https://en.wikipedia.org/wiki/File:Human_Kozak_context._Version_2).

This consensus allowed construction of an ATG context score based on agreement with the consensus and the information content of the position. The distribution of scores for reference initiators were contrasted to scores for 100,000 random sequences composed with nucleotide composition of the overall 5’UT-ome of *Chlamydomonas.* There was a clear separation between unimodal score peaks for reference initiator ATGs and the random control (Figure 4A).

**Figure 4.**
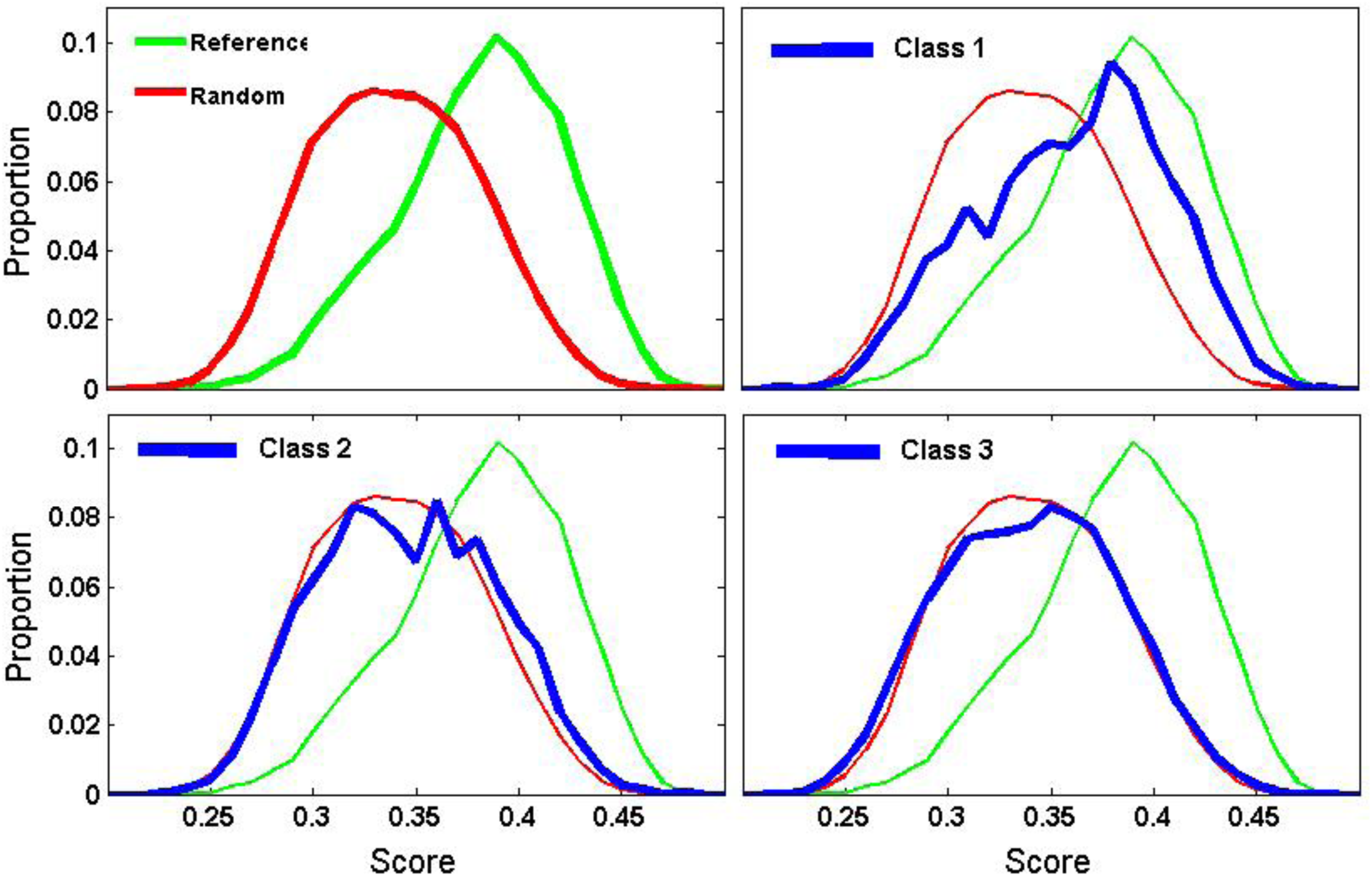
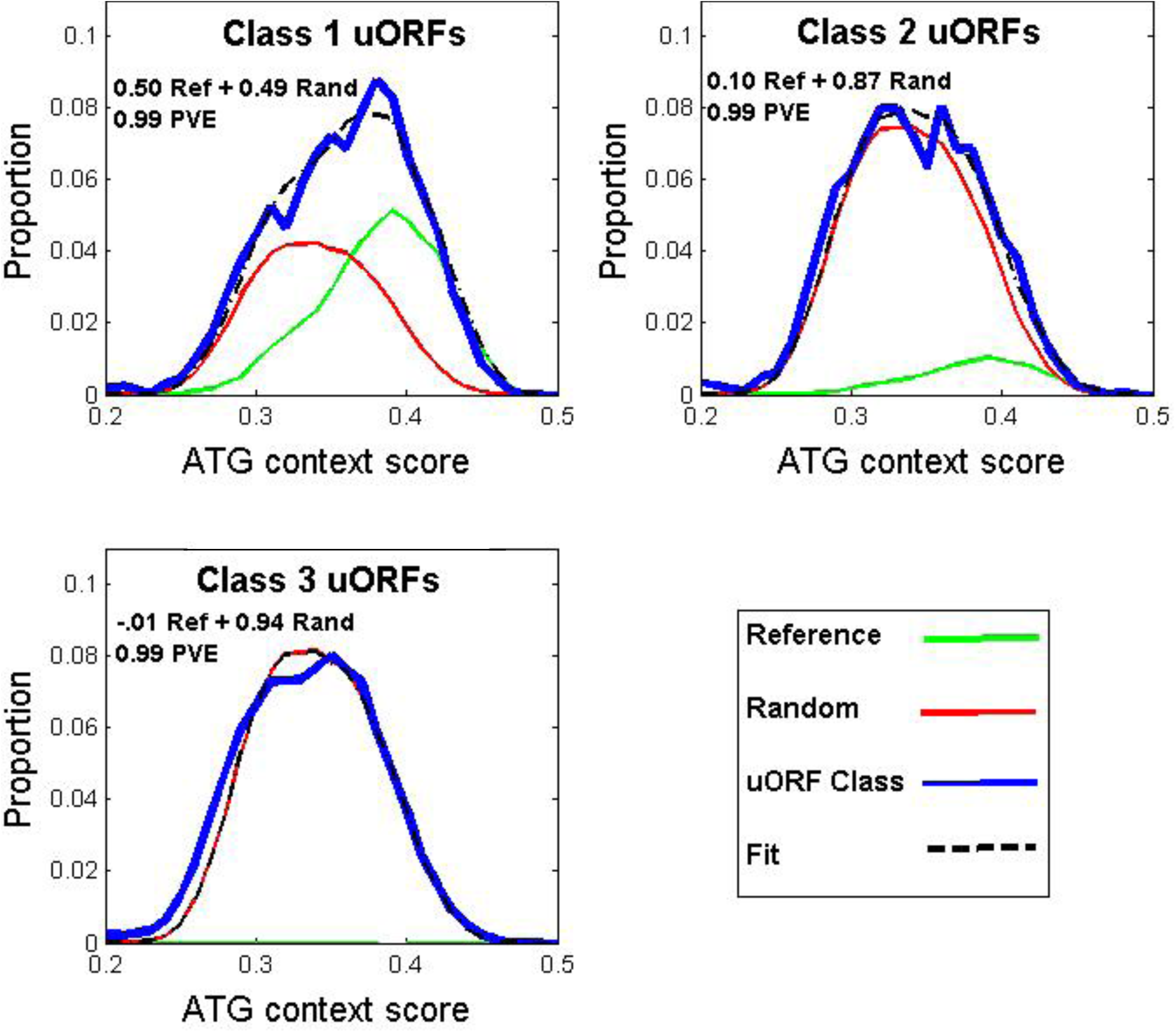
Class 1 uORFs, but not Class 2 or Class 3 uORFs, show significant agreement to the Kozak consensus. A: Reference, random and uORF agreement with the consensus. A ‘Kozak score’ was defined as the sum of bits corresponding to the observed nucleotides at each position surrounding the ATG (Figure 3). Top left: comparison of scores of reference initiator ATGs (green) to scores of 100,000 random sequences composed with the nucleotide frequency of 5UT (red). Remaining panels: uORFs (blue) compared to reference and random distributions. **B: Projection of uORFs onto space spanned by reference initiators and random sequence.** Define Matrix A = [distribution of reference;distribution of random]; vector C=distribution of uORF; then [x y]= (A^⊤^A) ^−1^ A^⊤^C gives the least squares best-fit solution for x ^*^ reference distribution + y ^*^ random distribution ~= C (Strang, 2009). The best fit is the dotted black line; weights and proportion of variance explained (PVE) are indicated.

### Distribution of Kozak consensus scores suggests that some Class 1 uORFs may be translated

Class 2 and Class 3 ATGs had a distribution of scores nearly identical to the random-sequence control (Figure 4A). Thus, if the Kozak consensus enhances initiation efficiency, many Class 1 and 2 ATGs may be skipped in favor of the downstream reference ATGs.

Class 1 ATGs had a very different distribution of Kozak consensus scores, which resembled a bimodal mixture of score distributions similar to the randomized control, and similar to the reference ATGs. A simple linear algebra calculation yielded the optimal mixture: a 51:49 combination of these distributions yielded a very good fit to the Class 3 distribution (99% of variance explained by this linear fit) (Figure 4B).

This observation suggests the hypothesis that Class 1 uORFs are heterogeneous. Half of them might be inefficiently translated, thus resembling the Class 2 and Class 3 uORFs. Half, on the other hand, might be translated either as alternative or as the exclusive *in vivo* initiations. Such initiation would result in a peptide with an N-terminal uORF fused to the reference peptide.

### Test of translation-dependent evolutionary selection on class 1 uORFs

If class 1 uORFs are translated and the resulting N-terminal extensions are under evolutionary constraint, then the N-terminal sequence could extend the alignment of the predicted peptide, when compared to other organisms. *Volvox* is a multicellular species with a recent common ancestor with *Chlamydomonas.* Many *Chlamydomonas* peptides have highly similar orthologs in *Volvox* (Prochnik et al., 2010). However, neutral nucleotide sequence divergence between *Volvox* species with and *Chlamydomonas* is >50%, based on substitution rates at neutral positions in highly conserved (thus alignable) proteins (actin, tubulin, CDKB). This level of sequence divergence means that sequence not under selection for its protein coding potential should rapidly lose any recognizable BLASTP (protein) homology, due to divergence and especially to fragmentation from gain of termination codons and loss of potential initiator ATGs. In contrast, if a sequence is translated and the peptide product under selection, then BLASTP homology will be retained. Figure 2A showed that a 50% divergence rate was equivalent to full randomization for completely eliminating enhanced lengths of class 1 uORFs.

The test was comparison of scores from BLASTP of the *Volvox* proteome against *Chlamydomonas* reference peptides, reference peptides with class 1 uORF N-terminal extensions, and two controls of reference peptides with class 1 uORF extensions that were scrambled at the level of predicted peptide. This scrambled control takes into account possible BLASTP score improvement due to simple-sequence features (e.g., poly-Pro aligning similarly with and without scrambling with Pro-rich N-termini in subject proteins). The test statistic was the maximum BLASTP score due to the uORF extension compared to scrambled controls. (This score is in units of bits, which are logarithmic and additive; thus the arithmetical difference in scores is an appropriate indicator of differential effect of the uORF). In many cases the real uORFs, but not the scrambled controls, increased the score by up to many hundreds of bits. Quantitative comparison of cumulative results suggests that at least 25-30% of class 1 uORFs are under selection for translated sequence content (Figure 5). This is likely a lower bound for the proportion of these class 1 uORFs that are included in *in vivo* peptides: first, because many proteins lack BLASTP homology at their N-termini, whether extended or not; second, because the comparable stretch in the *Volvox* annotation might have *also* been incorrectly assigned to an untranslated uORF.

**Figure 5.**
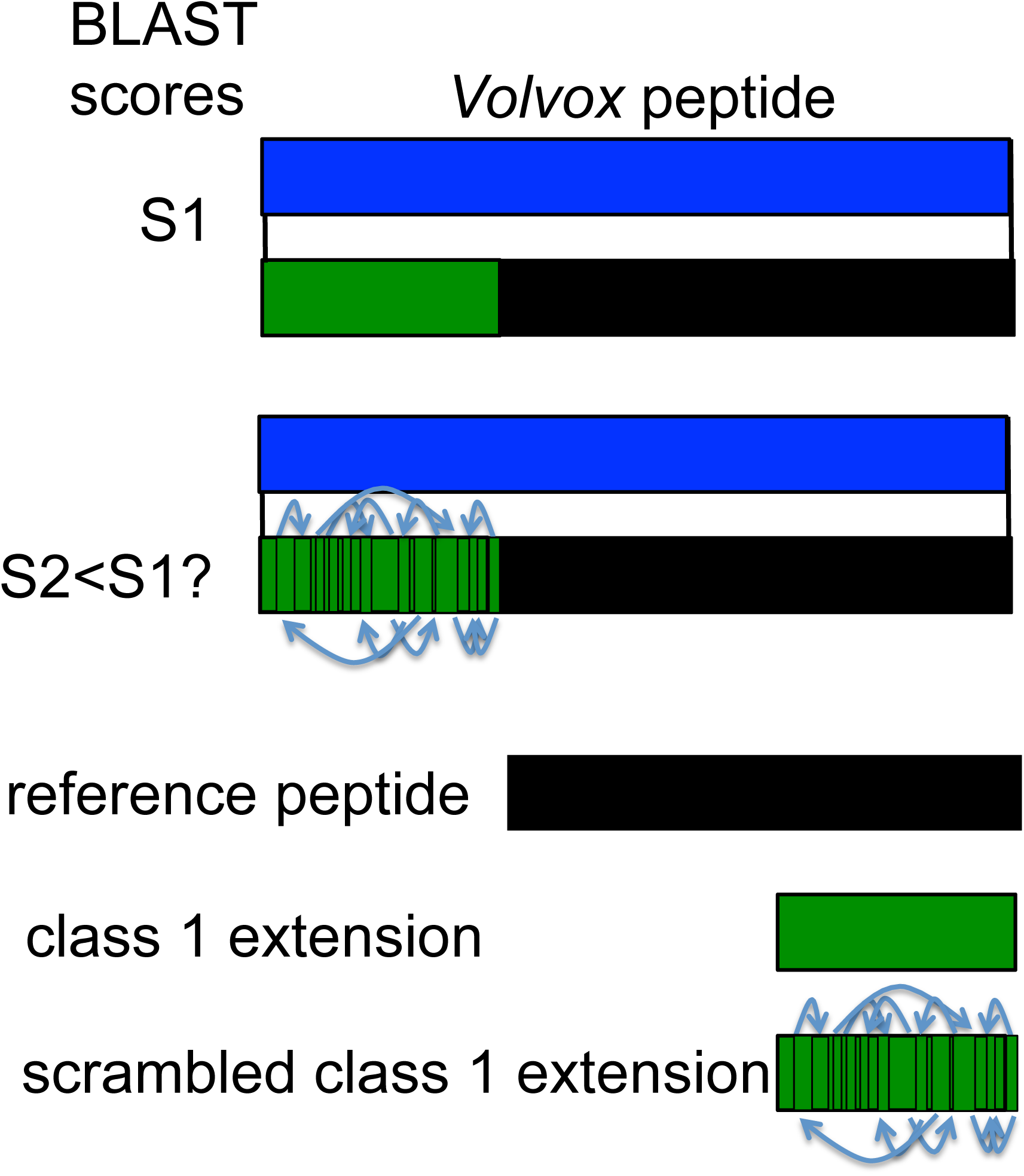
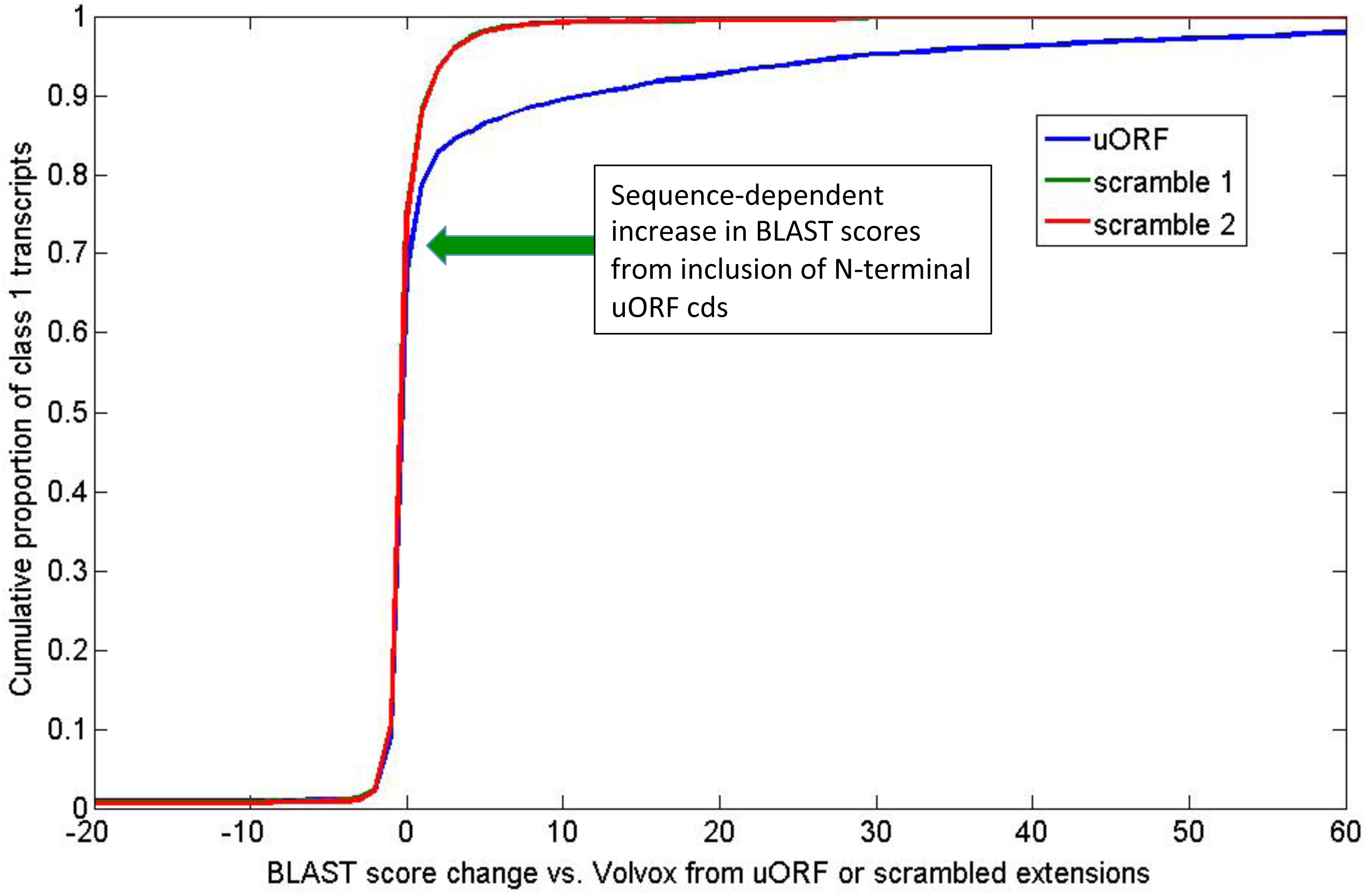
Many class 1 uORFs encode evolutionarily conserved N-terminal extensions. **A.** Schematic of the test. BLASTP alignments to the Volvox proteome were carried out for each class 1 transcript using four different versions: the reference transcript; the reference N-terminally extended by the class 1 uORF; and the reference transcript N-terminally extended by scrambled versions of the class 1 uORF peptide. Sequence-dependent BLASTP score improvement (S1>S2, S2 S3) was taken to argue for evolutionary conservation of class 1 coding sequence. **B.** The differences between maximal BLASTP scores of the reference peptide and the N-terminally extended versions (class 1 uORF peptide, or the scrambled uORF peptide for all class 1 transcripts is plotted as cumulative distributions. The divergence of the uORF peptide from the scrambled versions at a score of >4 and about 70-75% of transcripts indicates sequence-dependent score increase (that is, score increase specific to the uORF and not to scrambled versions) in 25-30% of class 1 transcripts.

A few examples of the aligned sequences resulting from these BLASTP comparisons are presented in Figure 6. It is clear in these cases that the uORF encodes evolutionarily conserved sequence, relevant to the function of the peptide. The data comprise a continuous series from such obvious cases to addition of only a few amino acids, with marginal or no effect on BLASTP scores.

**Figure 6.**
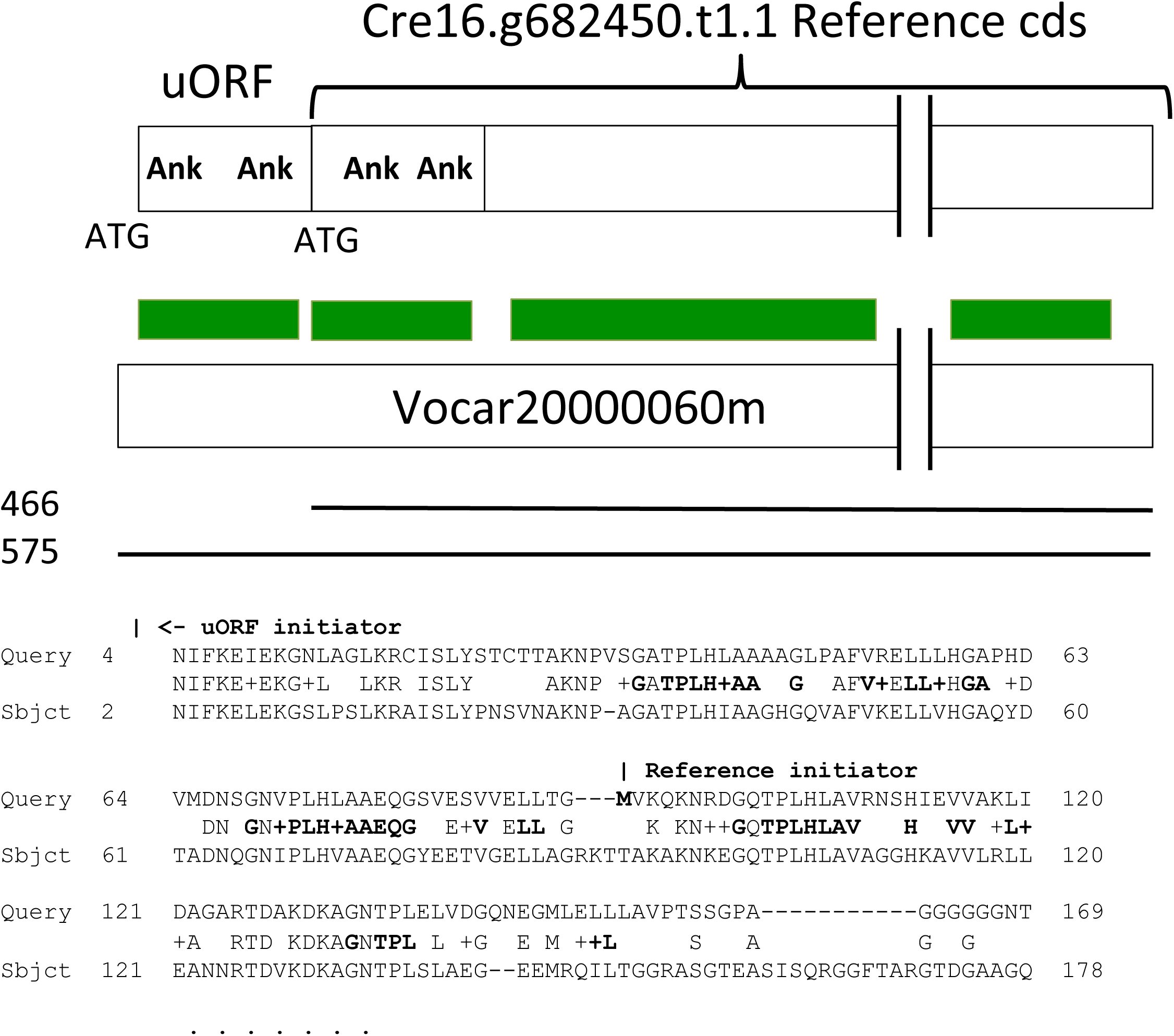
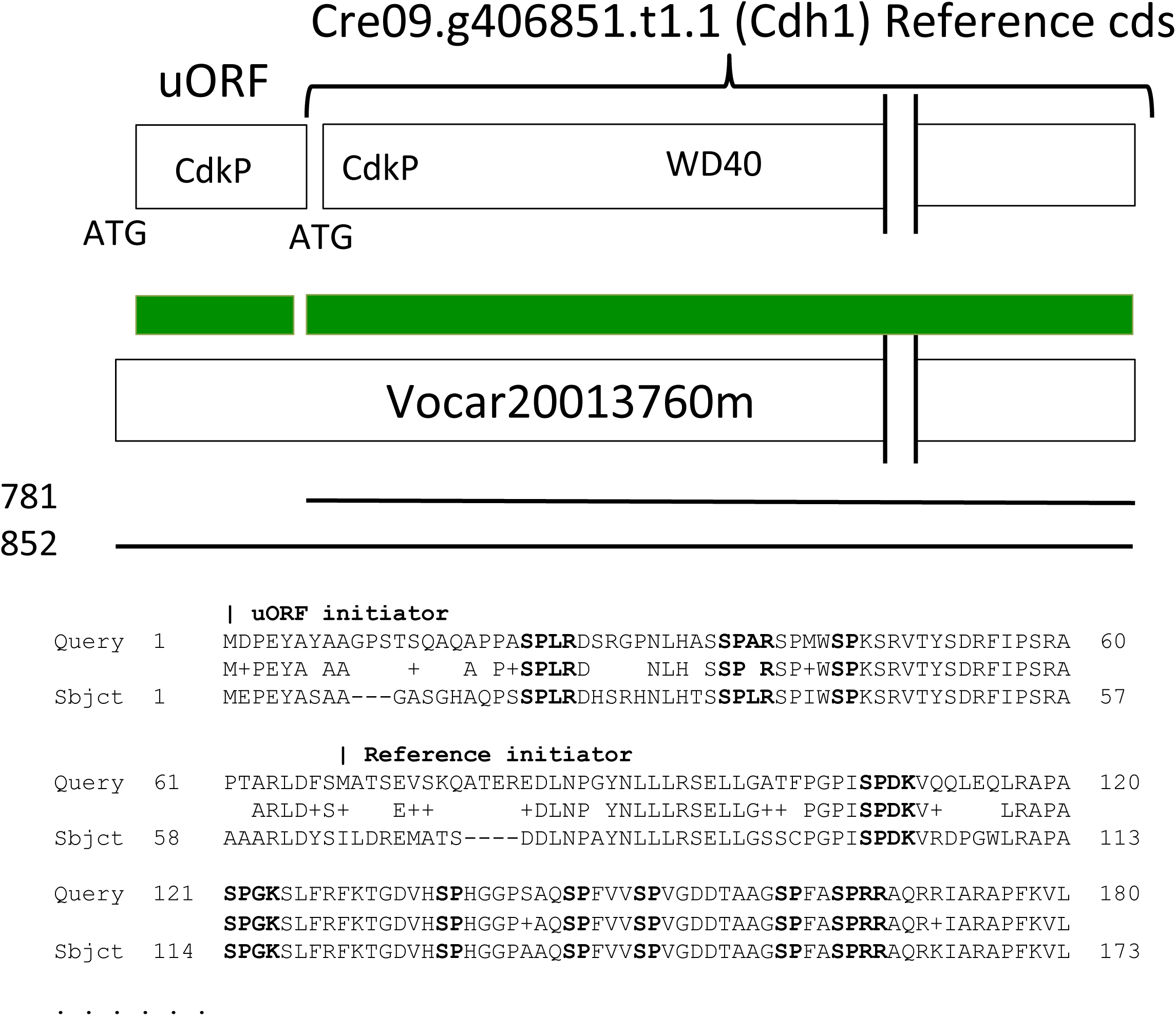
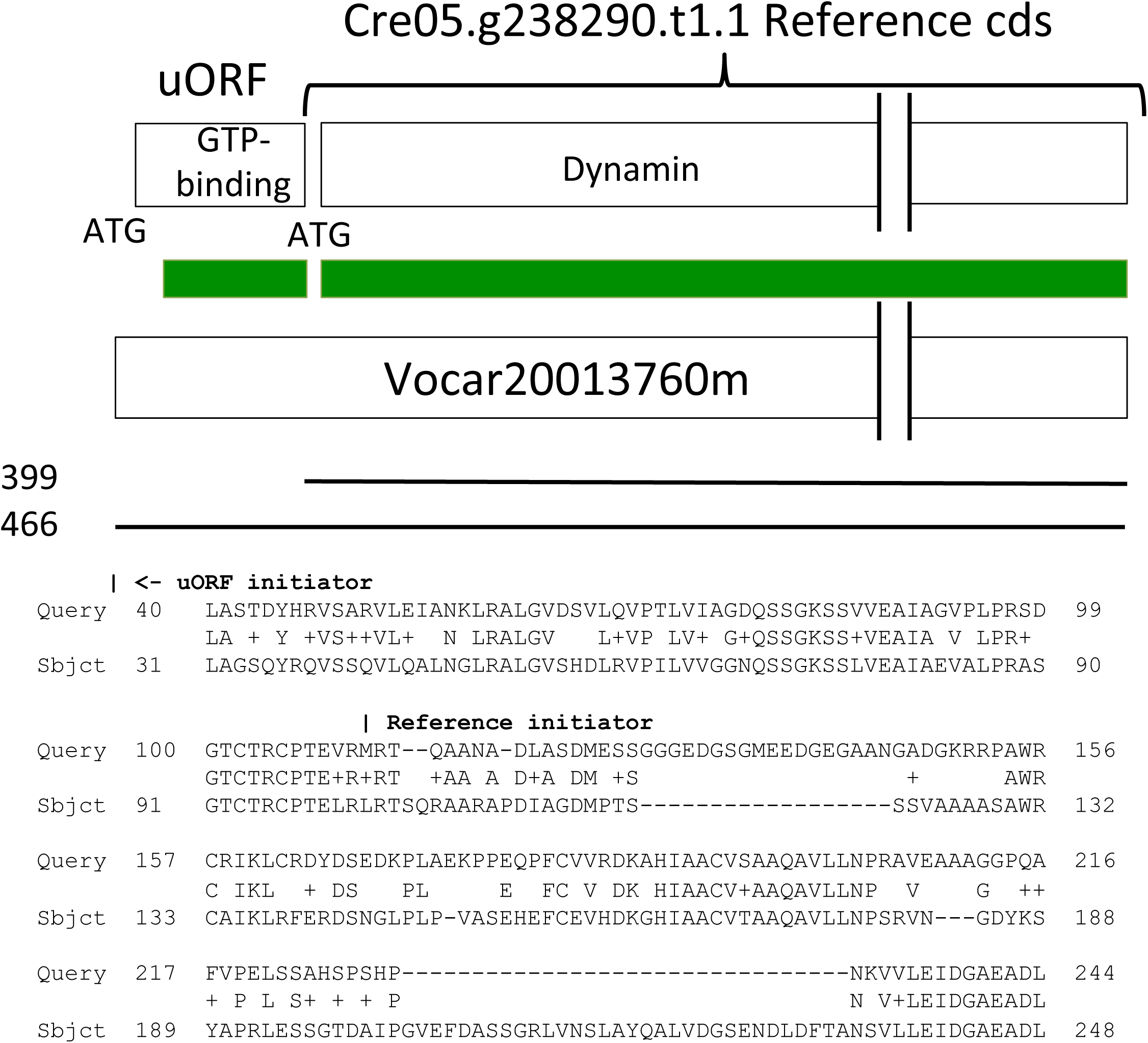
Examples of BLASTP score improvement by class 1 uORF N-terminal extensions. The N-terminal sequence of the best *Volvox* BLASTP hit to the reference coding sequence is shown, starting from the indicated ‘Reference Initiator’, along with the extended alignment from the class 1 uORF starting at the ‘uORF initiator. Regions of alignment are sketched in green and BLASTP scores indicated at left. **A.** Ankyrin repeat-containing protein. Two ankyrin repeats (bold in alignment below) are found in the N-terminal extension, and two more in the reference coding sequence. **B.** Cdh1. Cdh1 is known to be regulated by cyclin-dependent-kinase phosphorylation (Zachariae et al., 1998). (minimal consensus S/T-P; extended consensus S/T-P-x-R/K). 3/7 such sites are in the class 1 uORF N-terminal extension. **C.** Dynamin-homologous protein, with characteristic GTP-binding domain of dynamins encoded in the class 1 uORF extension.

A complete tabulation of the results of the BLASTP analysis is provided in Supplementary Information Table 1.

### Correlation of high Kozak consensus score and probable *in vivo* translation for Class 1 uORFs

An apparent bimodal distribution of Kozak consensus scores among Class 1 uORFs led to the suggestion above that the higher-scoring Class 1 uORFs might be preferentially translated. Figure 7A shows Kozak score distribution of the Class 1 ATG and the reference ATG from the upper ~25% of BLASTP improvement, compared to the remainder. The high-scoring cases had Kozak scores indistinguishable from overall reference ATG initiators, and interestingly had significantly better scores than the reference initiators from the same set of transcripts. The lower 75% cases were essentially indistinguishable from the bulk case (compare Figure 7A to Figure 4A, top right).

**Figure 7.**
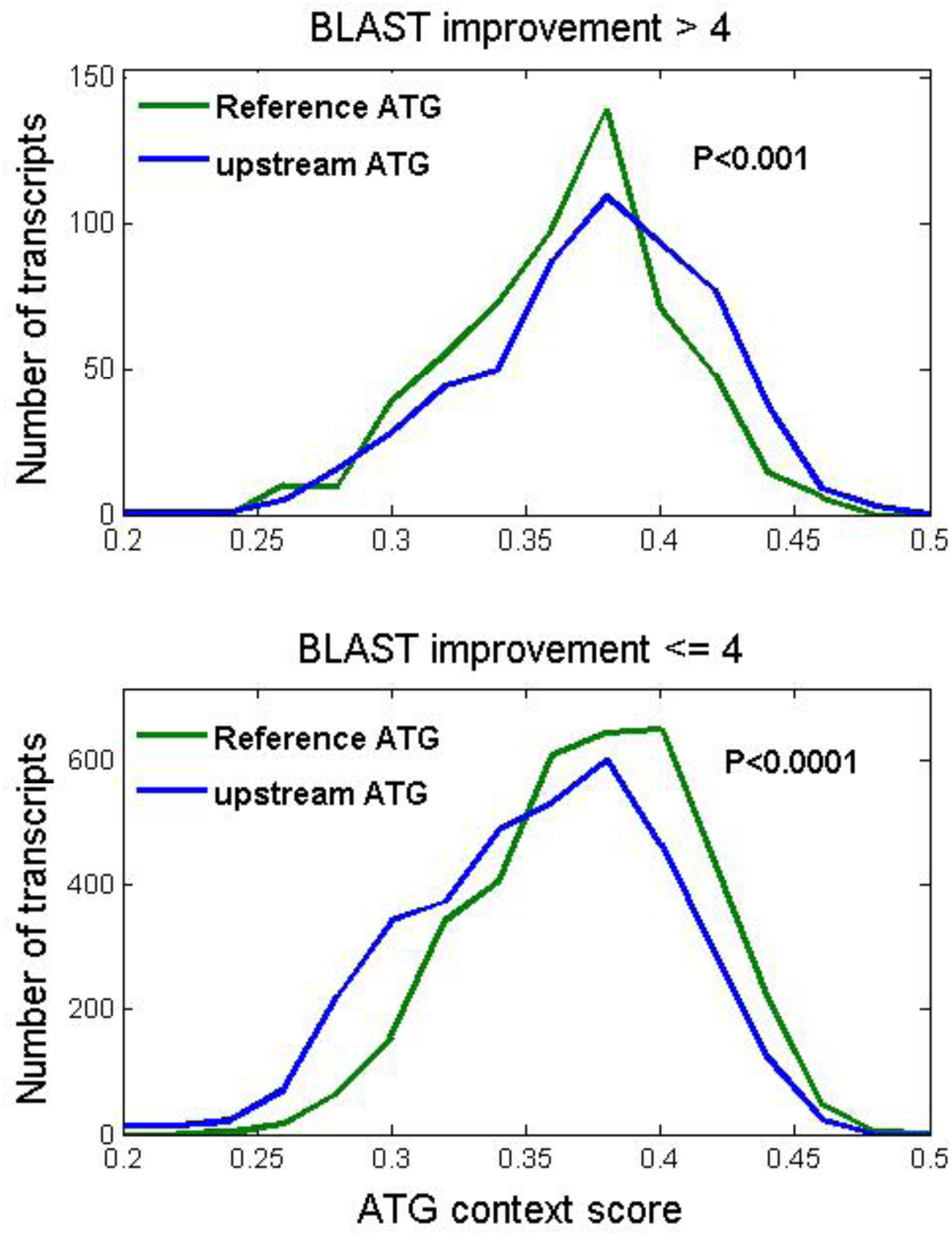
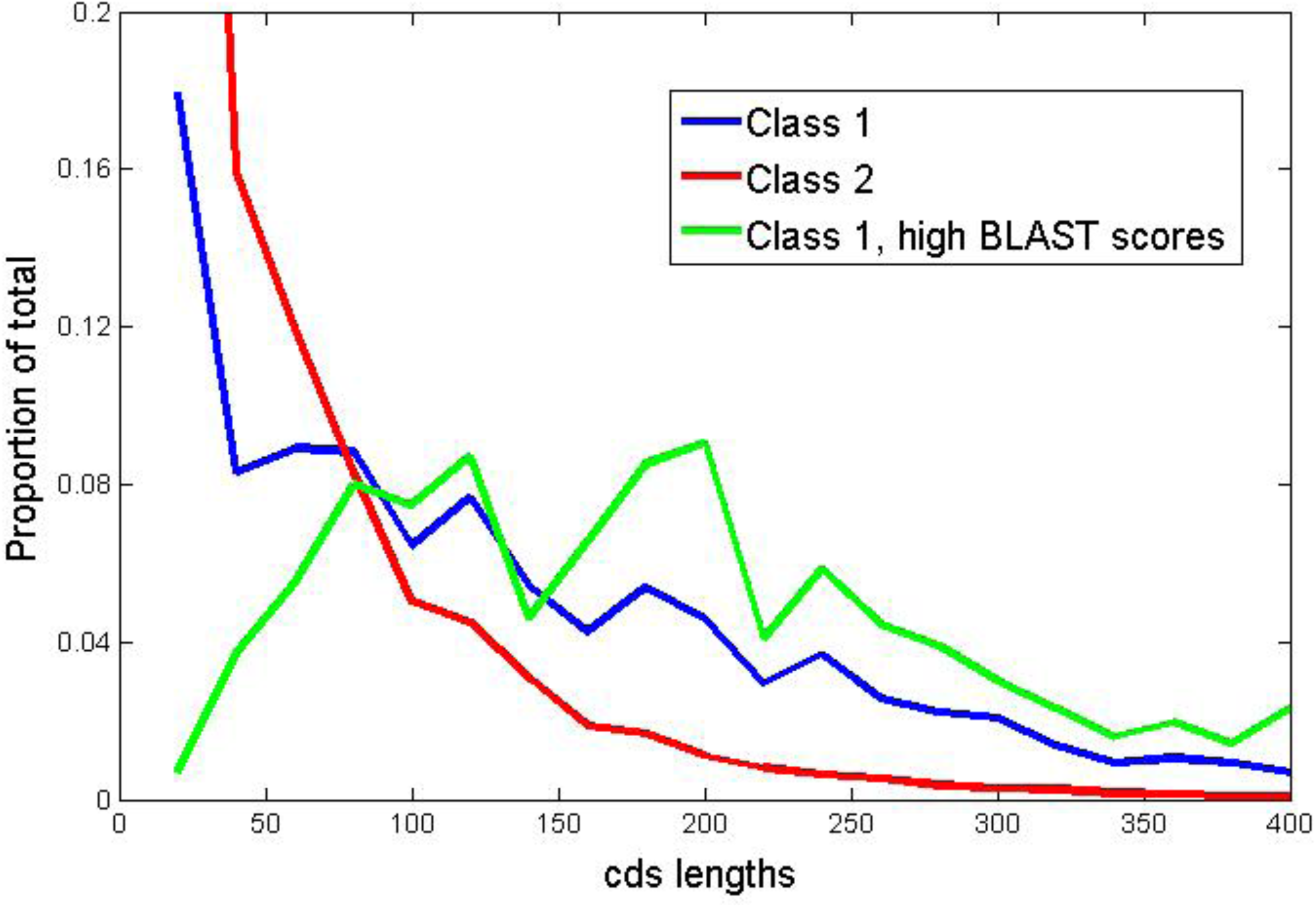
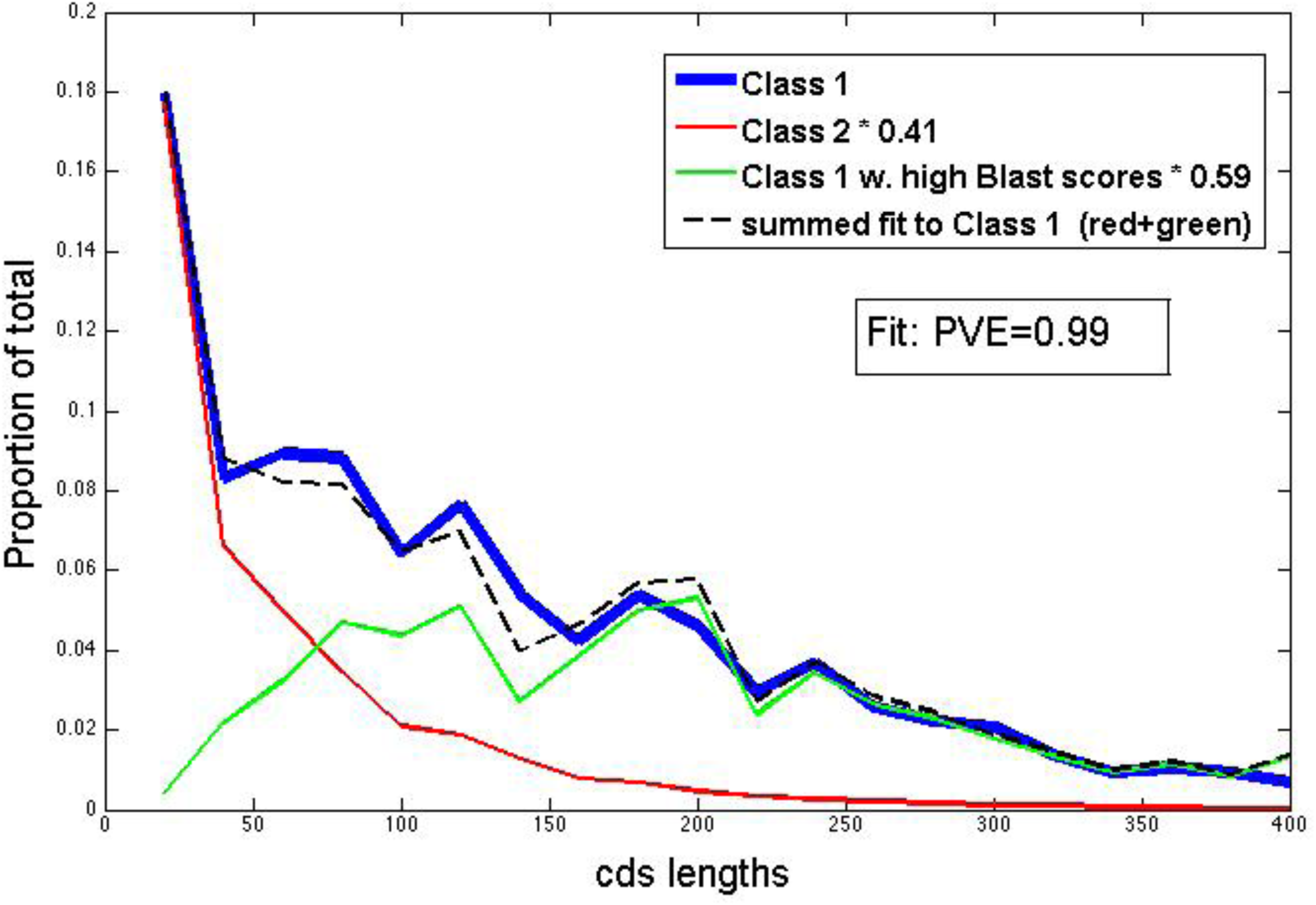
Class 1 uORFs with high BLASTP scores have high ‘Kozak’ scores and long coding sequences. **A.** Above: Kozak scores for the class 1 uORF ATG and the reference ATG, for all transcripts in the high BLASTP-improvement score class (see Figure 5). Below: Kozak scores for the same ATGs in the low BLASTP-improvement score class. Insets: probability by t-test in the high BLASTP class that reference ATGs have average scores greater than or equal to uORF ATGs (above), probability in the low BLASTP class that reference ATGs have average scores less than or equal to uORF ATGs (below). **B.** High BLASTP-score class 1 uORFs are longer than average class 1 uORFs. **C.** The length distribution of the total pool of Class 1 uORFs can be accounted for as a 0.59:0.41 sum of the high BLASTP-score subset distribution and the Class 2 uORF distribution, by least squares best fit as in Figure 4B.

This finding supports the idea that the Kozak consensus is relevant to translation efficiency, and the idea that a substantial proportion of class 1 uORFs are translated *in vivo,* either as alternative or as the sole initiation codons.

Class 1 uORFs are significantly longer on average than either class 2 uORFs, or class 1 uORFs from ‘mutagenized’ or randomized sequence (see above). This length effect was markedly enhanced for the class 1 uORFs in the upper 25% of BLASTP improvement (Figure 7B). If class 1 uORFs are not in fact translated, there is no obvious reason for any length difference compared to class 2 uORFs, since they differ only in translational frame relative to the reference (Figure 1). The total class 1 cds length distribution could be modeled as a sum of the class 2 distribution and the high-BLASTP-scoring class 1 distribution with ~40:60 mixture (Figure 7C). This split is similar to the 49:51 split in the fit to Kozak consensus scores; both results suggest that around half of class 1 uORFs are translated efficiently. (Note that if the class 1 uORFs are a heterogeneous 50:50 mixture of neutral and functional sequences, then considering only the neutral class, the lengths and sensitivity to mutagenesis and randomization becomes very similar to class 2; and their numbers become just about half that of the class 2 uORFs, as expected given three reading frames).

### Weighted predictions for translation of class 1 uORFs

The results above suggest the likelihood that approximately 50% of class 1 uORFs are alternative or exclusive sites of *in vivo* initiation. In cases where a strong improvement in BLASTP score could be detected by including the uORF N-terminal extension, these transcripts can be identified directly (Figures 5, 6). Such cases are a minority, though, and it would be desirable to have a sense of the likelihood of translation initiation for the entire class of ~4000 class 1 uORFs.

Class 1 uORFs that are more likely to be translated have two sequence features (independent of *Volvox* alignments): the uORF coding sequence is longer, and the Kozak consensus is stronger. Neither feature is quantitatively strong enough to form a digital classifier. The two-dimensional differentiation between uORF classes 1, 2, and 3, and the ‘high BLASTP’ subclass of uORF class 1 was a stronger separator (Figure 8A). The surface for class 1 could be almost exactly modeled as a 50:50 split between the class 1-high BLASTP subclass (representing efficiently translated uORFs) and class 2 (representing presumably poorly translated uORFs) (Figure 8B). This allowed construction of a simple Bayesian test for the likelihood that a class 1 uORF is translated based on its coding sequence length and its Kozak score, using as training sets the high-BLASTP class 1 subset (positive examples) and class 2 uORFs (negative examples). The strength of discrimination is illustrated in Figure 8C; the receiver-operator characteristic (ROC) curve shows the false positive/true positive relationship for various probability cutoffs. While this test is still certainly not definitive in any individual case, it does provide an empirically based estimate of the probability of translation in the complete set of class 1 uORFs (Supplementary Table 1). In advance of more definitive empirical evidence, these probabilistic classifications may be helpful as an adjunct to the (necessarily) digital summary in Phytozome that some sequence ‘is’ or ‘is not’ part of the coding sequence.

**Figure 8.**
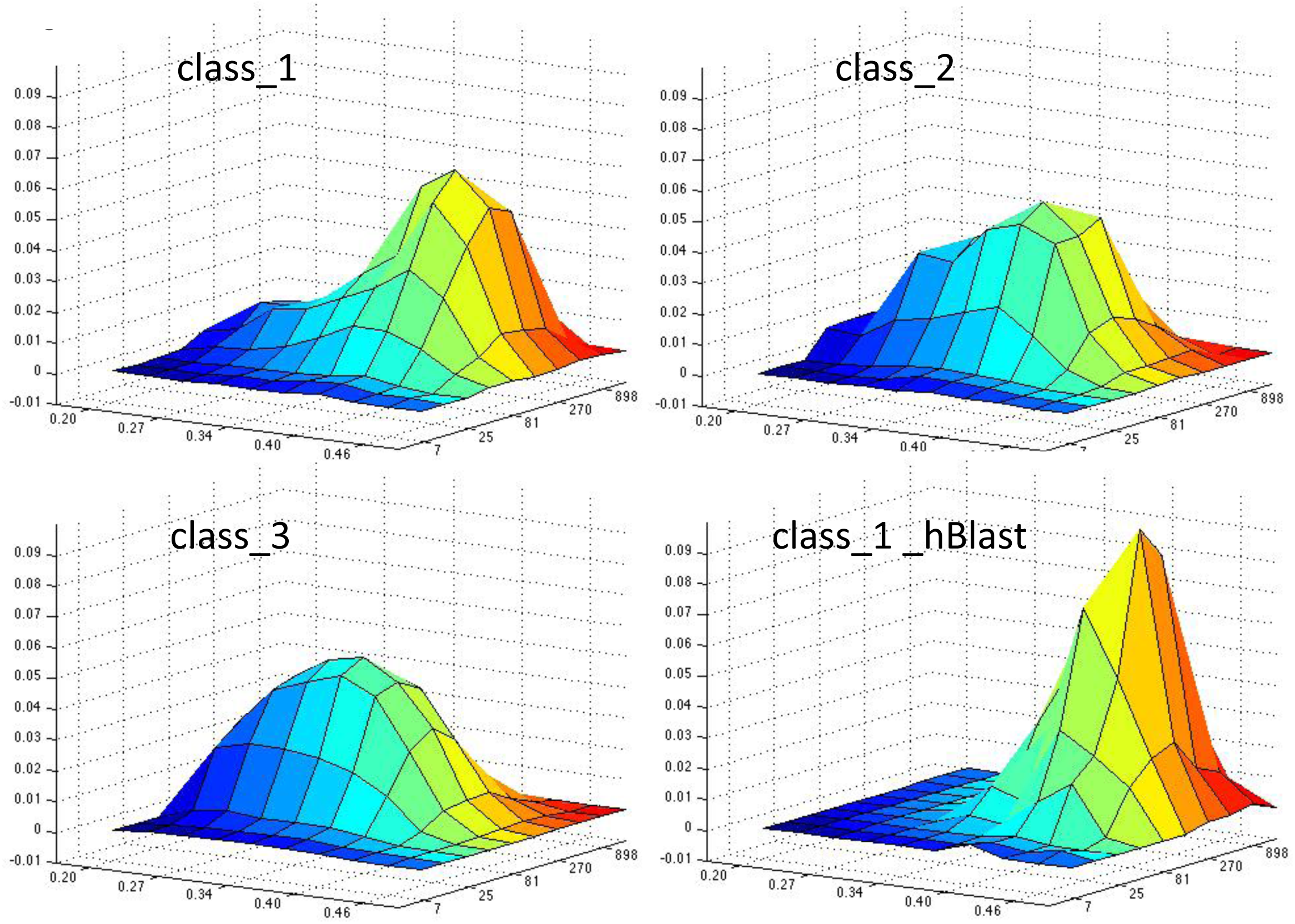
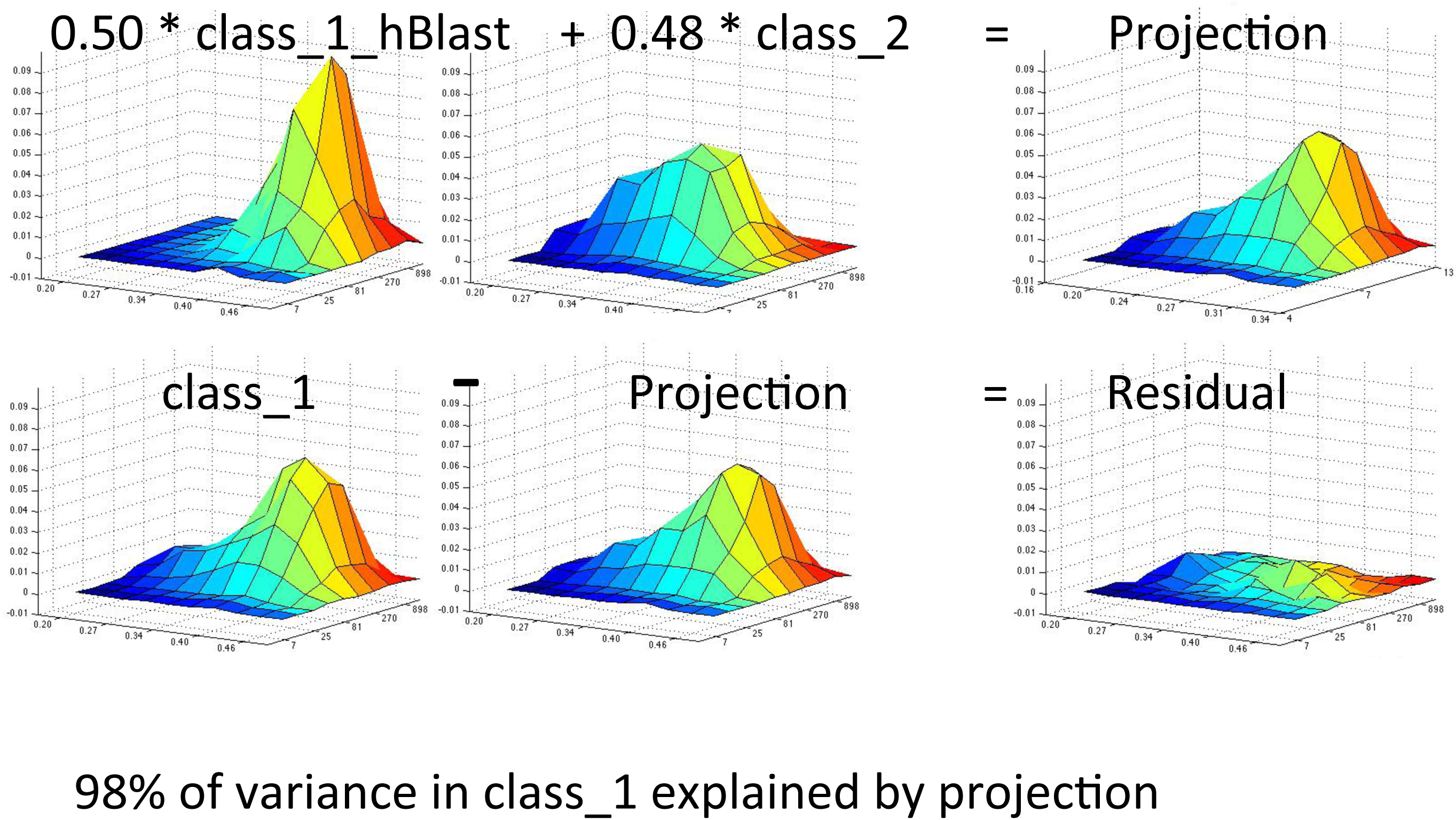
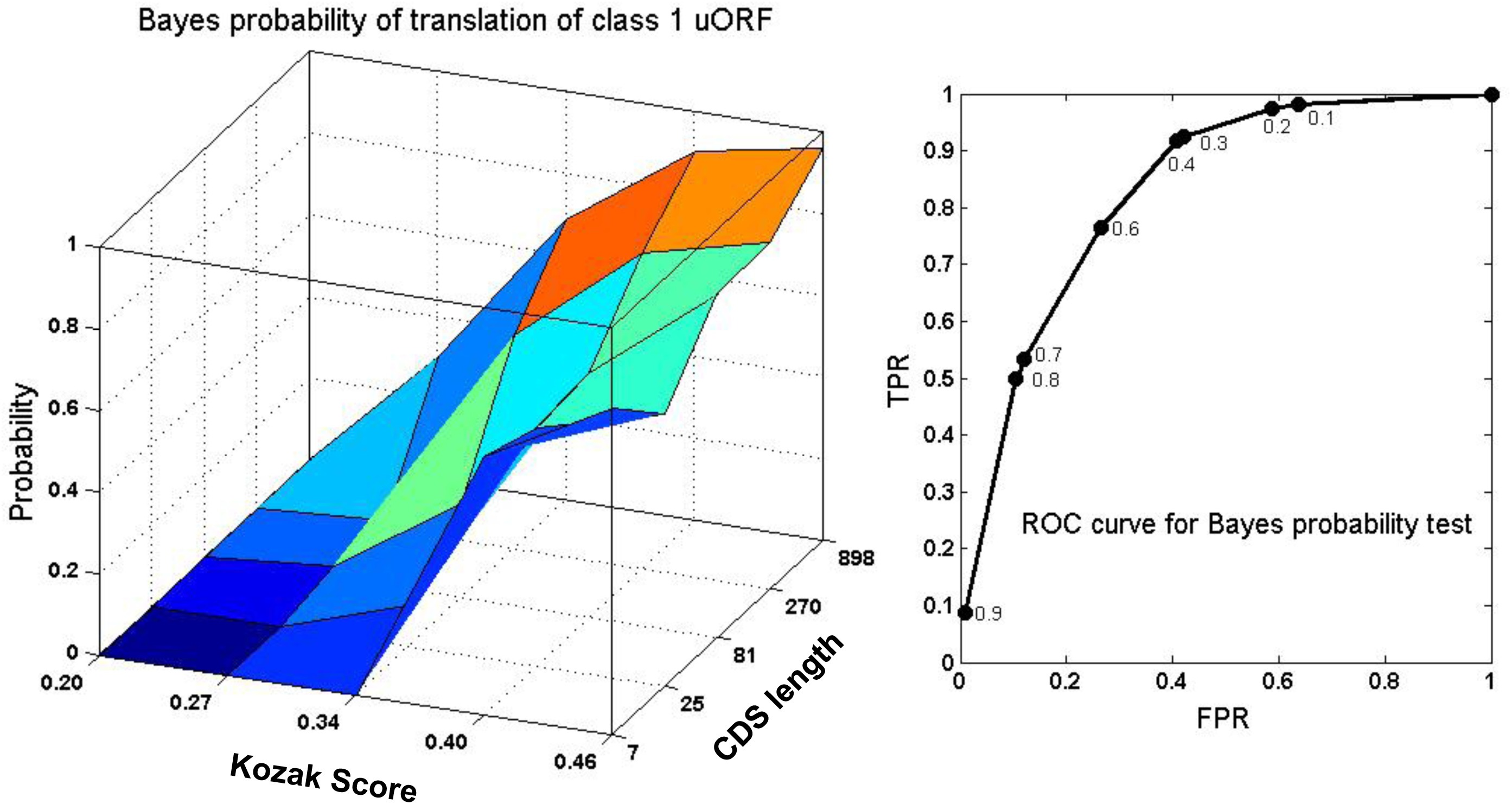
Two-dimensional accounting for uORF variability. **A.** Surface of proportion of uORFs over a two-dimensional grid of Kozak score (exponentiated) and log coding sequence length. These transformations were chosen because the Kozak score is based on an essentially logarithmic information scale (Figure 3), whereas coding sequence lengths can be conceptualized as due to exponential decay from finite probability of hitting a stop codon (class 3) or the beginning of the coding sequence (classes 1 and 2). Classes 2 and 3 have broad peaks with low Kozak scores (left axis) and shorter coding sequences (right axis). The high BLASTP-score subclass of class 1 uORFs (lower right) has a sharp peak at high Kozak score and longer coding sequences. The complete pool of class 1 uORFs appears heterogeneous, with a peak similar to the high BLASTP subset and a shoulder similar to the distributions of classes 2 and 3. **B.** Least-squares accounting for the two-dimensional class 1 uORF distribution as a sum of 0.50 high BLASTP class 1 uORF subset and 0.48 class 2 uORF (matrix calculation as in Figure 4B); 99% of variation is explained. **C.** A Bayesian test for translation of class 1 uORFs. Assuming ~50:50 split of translated and untranslated class 1 uORFs (Figures 4C, 7C, 8B), then Bayes’ theorem simplifies to P(translated | Kozak score & cds length)=P(Kozak score & cds length | translated) / (P(Kozak score & cds length | translated) + P(Kozak score & cds length | not translated). We assume that P(Kozak score & cds length | translated)=P(Kozak score &cds length | member of high-BLASTP class), and P(Kozak score &cds length | not translated)= P(Kozak score &cds length | class 2 uORF), based on Figures 7, 8A, B. Then P(translated | Kozak score &cds length)= proportion(high-BLASTP-score class 1 uORF with this Kozak score & cds length) / (proportion(high-BLASTP-score class 1 uORF with this Kozak score & cds length) + proportion(class 2 uORF with this Kozak score & cds length)). This surface is graphed at left. At right is the receiver-operator characteristic (ROC) curve given this surface, for varying cutoffs of nominal Bayesian probability, with the assumption that all translated class 1 uORFs will have sequence properties similar to the high-BLASTP score subset, while all untranslated class 1 uORFs will be similar to (the presumably untranslated) class 2 uORFs.

The conclusion that a substantial proportion of class 1 uORFs is translated is based on multiple comparisons and tests, that are entirely independent: length distributions of class 1 cds compared to classes 2 and 3, and to randomized controls; Kozak score distributions of the same set, also compared to reference ATGs; BLASTP improvement by comparison to the *Volvox* proteome. The estimated 50:50 split is also based on projection of class 1 uORF data onto multiple distint subspaces: spanned by Kozak scores of reference ATGs and random sequence; and spanned by cds length of the high-BLASTP class 1 subset and class 2 uORFs, alone or in two-dimensional combination with Kozak scores.

### A large majority of reference ATGs are likely contained in coding sequence

The analysis so far was focused on the question of whether the reference ATG or a more 5’ ATG was a more likely site of translation initiation. A converse question can be asked: could some reference ATGs in fact *themselves* begin uORFs, with the authentic *in vivo* start codon being one annotated as internal? Another BLASTP comparison to *Volvox* provides insight. In this comparison, I first determined the subset of transcripts for which there was a detectable *Volvox* BLASTP hit for which the maximum score was dependent on sequences immediately 3’ to the first internal ATG after the reference ATG. There were 8320 such transcripts. In 78% of this set of transcripts, the segment between the reference ATG and the next ATG contributed further to the BLASTP score (Figure 9A), which would be unexpected if the ‘reference’ ATG was not the initiator. This could be taken to imply that as many as 22% of ‘reference’ ATGs are not, in reality, part of coding sequence. Arguing against this idea, though, Kozak scores were on average higher for the reference than for the first internal ATG (P<<<0.001 by t-test for both comparisons) independent of BLASTP results (Figure 9B). Therefore, at least 78%, and likely a higher proportion, of the reference ATGs are actually within coding sequence.

**Figure 9.**
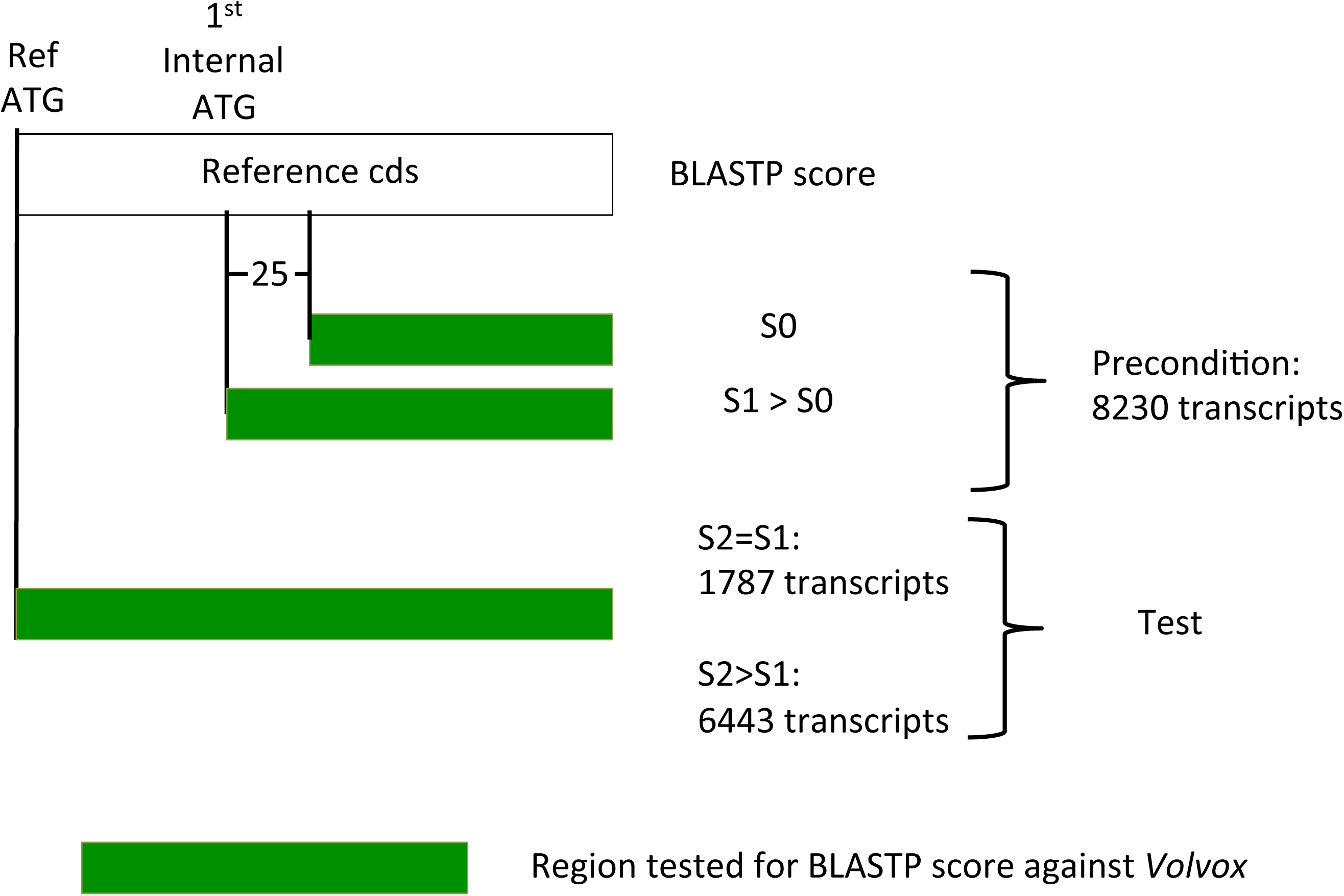
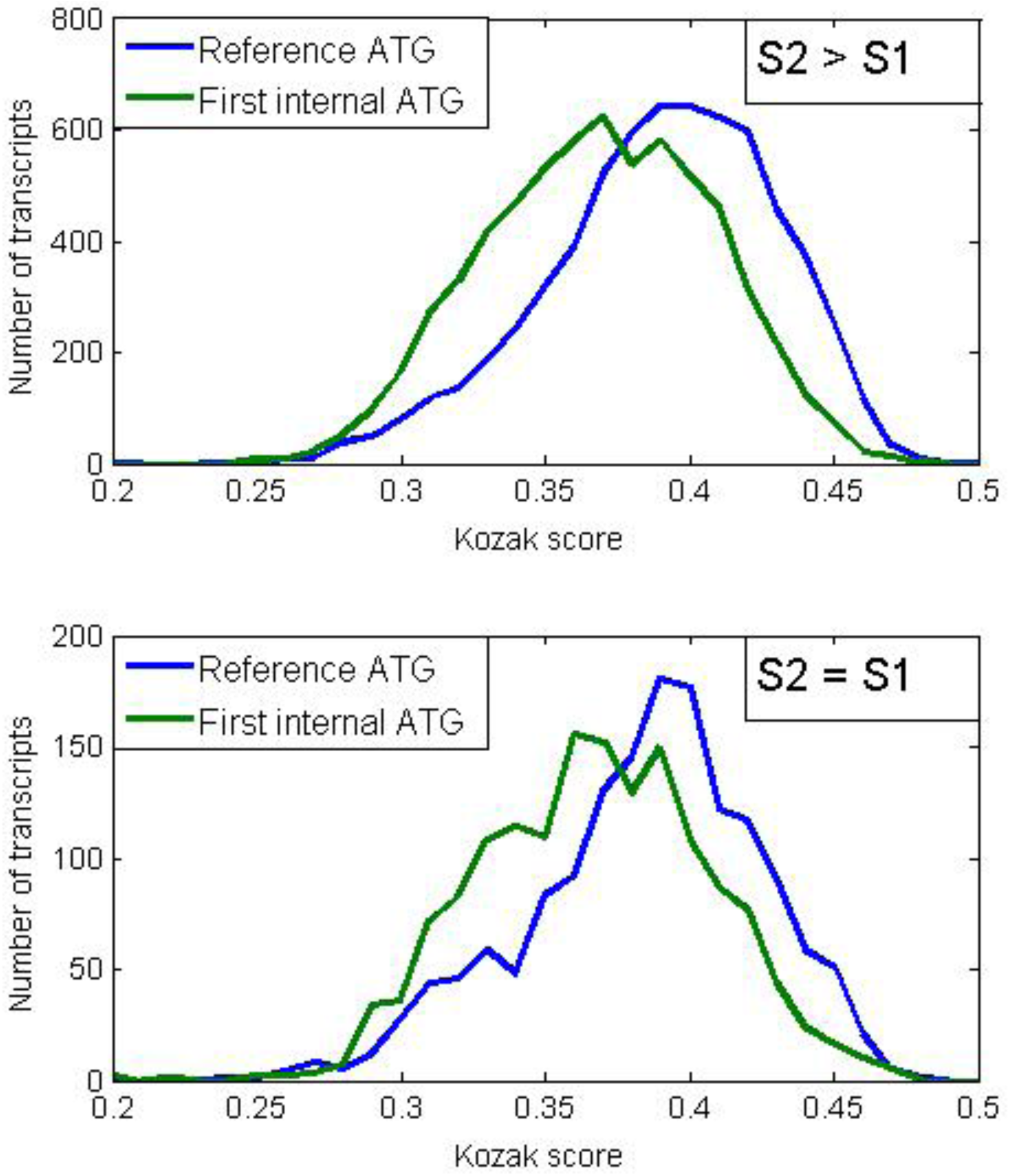
Test for reference ATG being in coding sequence. **A.** In many coding sequences there will be BLASTP homology to a *Volvox* peptide, starting alignment 25 amino acids downstream of the 1^st^ internal (post-reference) in-frame ATG, with score S0 (green bar indicates region of alignment). In a subset of these cases this alignment and BLASTP score will increase if the query for alignment is extended to start exactly at the 1^st^ internal ATG (score S1 > S0). This is the precondition for the test, which is met in 8230 transcripts. In such cases, if the reference ATG is the genuine initiator **or** 3’ of the genuine initiator, it is quite likely that the score will increase again to S2 > S1 if the BLASTP query additionally includes the segment encoded between the reference and second ATG. In contrast, if the ‘reference ATG’ is **not** translated, then the score will almost surely not increase (S2=S1), since conservation to *Volvox* should require selection based on continued translation to peptide. In 78% of transcripts fitting the precondition S1>S0, we observed S2>S1, setting an upper bound of 22% for the frequency of reference initiators not actually translated. **B.** Comparison of Kozak scores for the reference ATG to those for the first internal in-frame ATG, in the two cases indicated in part A. In both cases the reference ATG population had significantly higher Kozak scores than the first in-frame ATG, suggesting that the reference initiator is a more probable *in vivo* site of translation initiation even in the cases below failing the BLASTP score test.

### Downstream open reading frames

The high prevalence of class 3 uORFs in the annotation suggests the possibility of translation re-initiation, so that if a class 3 uORF is translated this does not block translation of the main reference coding sequence. If such reinitiation is common, it raises the possibility that open reading frames in the 3’ untranslated region (downstream ORFs or ‘dORFs’) might similarly be translated after termination of the reference coding sequence. There are abundant open reading frames in the annotated 3’-untranslated regions - ~300,000 in all (Figure 10, top left), mostly less than 100 nt in length. A randomized control has about half as many dORFs. The length distributions and distributions of Kozak scores are identical between the real and randomized data. These observations do not strongly suggest that dORFs are translated in great abundance, although the increased number compared to the randomized control suggests some structure to this population.

**Figure 10.**
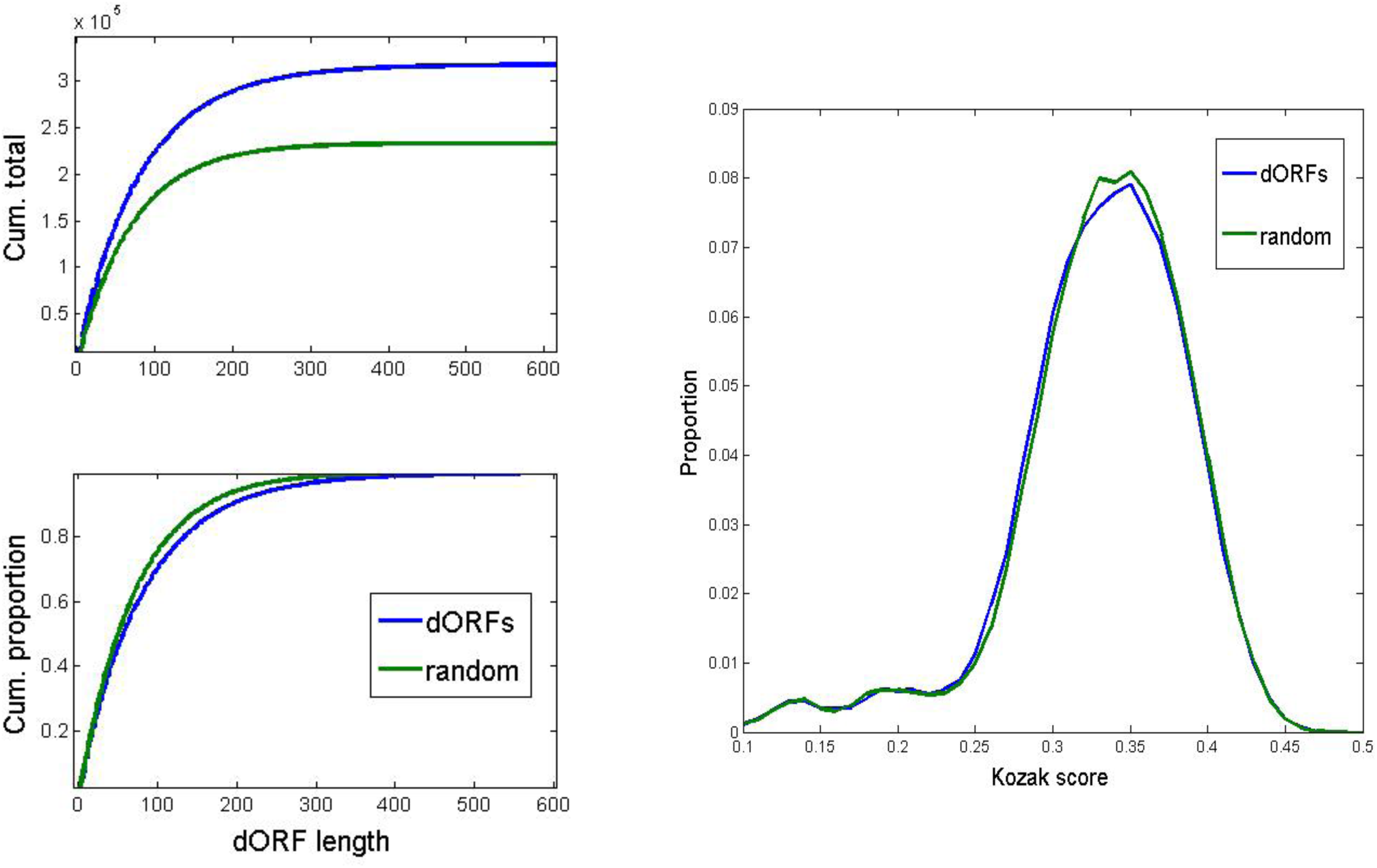
Downstream open reading frames. Open reading frames within annotated 3’ UTR sequences (downstream open reading frames or dORFs) were quantified with respect to length, and compared to uORFs from a randomized control with the same overall nucleotide composition and same length distribution. Top left: cumulative numbers; bottom left: cumulative proportions. Right: distribution of Kozak scores for real and random dORF ATGs.

## Discussion

### The translational fate(s) of abundant uORFs in the annotated *Chlamydomonas* genome

The annotated *Chlamydomonas* genome contains a very high number of uORFs in annotated 5’ untranslated sequence. Since a large majority of these uORFs are upstream of a reference coding sequence that is evolutionarily conserved, it is very likely that the uORFs do not fully block translation of this reference.

This leaves a small number of possibilities to explain the uORFs. (1) The transcription start site is incorrectly annotated in a majority of *Chlamydomonas* transcripts, and transcripts in fact generally initiate hundreds of nucleotides 3’ of the annotated position. This is logically possible; however, it is contradicted by EST evidence in the case of a substantial number of transcripts (Phytozome website). (2) The uORFs are not sites of translation initiation, due to sequence constraints (such as the Kozak consensus) resulting in their inefficient use. Our data support this possibility for classes 2 and 3 uORFs, and for about half of the class 1 uORFs. (3) The uORFs are sites of translation initiation, but do not interfere with translation of the reference cds. This is our interpretation for about half of the class 1 uORFs, which likely encode translated N-terminal extensions to the reference cds. It is an interesting possibility that class 3 uORFs are translated. Such translation can be compatible with translation of downstream AUGs (as in GCN4; Hinnebusch 2011); the GCN4 case also shows clearly that this situation can allow for regulated translation initiation. (Class 2 uORFs are likely not suitable for this mechanism, assuming that scanning is largely unidirectional). How widespread such translational control may be in *Chlamydomonas* is a question for the future; one case has been documented (Moseley et al., 2002).

### Functional consequences of N-terminal extensions

The class 1 uORFs vary tremendously in length, and it is likely that cases where only a few residues are missed at the N-terminus in the reference annotation will frequently have rather minor functional consequences. Also, protein N- and C-termini are probably more likely to be poorly folded than other regions (perhaps explaining why terminal epitope tagging so frequently is permissive for protein function). There are many cases, though, where functional consequences from exclusion of the class 1 uORF N-terminal extension are likely quite substantial (a few clear examples, selected from many, are in Fig. 6; it is also quite common for protein N-termini to contain relatively unstructured ‘addressing’ sequences for post-translational modification and/or subcellular localization, such as signal sequences for secretion or organellar transport). A different kind of consequence comes in searching for causative mutations following random mutagenesis: even a small, potentially unstructured N-terminal extension due to a uORF can be the site of a chain-terminating null mutation, which can go completely unrecognized if the uORF is not annotated as a possible contributor to coding sequence. In fact, we found just such a case in a screen for latrunculin B-sensitive mutants: a one-nucleotide deletion in an annotated 5’UT region resulted in a strong mutation in a specific molecularly identified complementation group (FC, M. Onishi, J. Pringle, in preparation). This finding was a major motivation for carrying out the present study; the deletion is, in fact, in a class 1 uORF. This ORF has a calculated probability of being translated of 0.90 by the Bayesian test described above (Figure 8C). Thus, this mutation is very likely an early chain terminator, consistent with other genetic results indicating that it produces a null allele.

### Genome annotation is probabilistic

It is a subtle problem that a sophisticated data presentation such as the *Chlamydomonas* annotated genome (Blaby et al., 2014) necessarily requires selection of a single one of a large number of alternatives; the most probable on some accounting is presumably selected, and is then represented ‘as if’ it were certainly correct – that is, every transcript model has one and only one translational start codon, the one indicated. However, the reasoning behind producing the object will most certainly be probabilistic and based substantially on statistical evidence. This was indeed the case for assignment of reference initiator ATGs in the *Chlamydomonas* annotation (M. Stanke, personal communication). A more accurate (though pedantic) description than ‘finding needles in a haystack’ (Blaby et al., 2014) might be ‘finding the most probable needles in a haystack that surely contains at least some’. An annotated genome fundamentally consists of a vast number of assertions that are statistically of the form (changing metaphors) ‘the first card dealt from the deck will not be the ace or king of hearts’ – usually but not always correct.

While this detailed examination of uORFs did uncover some issues and problems, it is likely that for the most part, the reference initiating ATGs are the authentic *in vivo* initiators. This is fortunate, and it is encouraging that a largely computational assignment of the complex biochemical events of mRNA production and translation can be correct in a large majority of cases. The explicitly probabilistic evaluation of likely sites of *in vivo* initiation presented here may be useful in consideration of the *Chlamydomonas* proteome, at least until there are sufficient data from proteomics and other approaches to settle the matter definitively.

## Acknowledgments

Funding support was from PHS 5RO1-GM078153. Many thanks to Masayuki Onishi for discussions of both uORFs and dORFs, and for finding the first examples of class 1 uORFs that motivated the broader study. Thanks to Sabeeha Merchant, Simon Prochnik and Mario Stanke for discussions and for clarifying aspects of gene annotation used in developing the *Chlamydomonas* reference. Thanks to Frej Tulin for extensive discussions and for carrying out BLAST searches.

## References

Altschul SF, Gish W, Miller W, Myers EW, Lipman DJ. Basic local alignment search tool. J Mol Biol. 1990 Oct 5;215(3):403–10.

Blaby IK, Blaby-Haas CE, Tourasse N, Hom EF, Lopez D, Aksoy M, Grossman A, Umen J, Dutcher S, Porter M, King S, Witman GB, Stanke M, Harris EH, Goodstein D, Grimwood J, Schmutz J, Vallon O, Merchant SS, Prochnik S. (2014). The Chlamydomonas genome project: a decade on. Trends Plant Sci. 2014 Oct;19(10):672–80.

Crooks GE, Hon G, Chandonia JM, Brenner SE. (2004) WebLogo: a sequence logo generator. Genome Res. 2004 Jun;14(6):1188–90.

Hinnebusch AG.Molecular mechanism of scanning and start codon selection in eukaryotes. Microbiol Mol Biol Rev. 2011 Sep;75(3):434–67

Kozak M. The scanning model for translation: an update. J Cell Biol. 1989 Feb;108(2):229–41.

Merchant SS, Prochnik SE, Vallon O, Harris EH, Karpowicz SJ, Witman GB, Terry A, Salamov A, Fritz-Laylin LK, Maréchal-Drouard L, Marshall WF, Qu LH, Nelson DR, Sanderfoot AA, Spalding MH, Kapitonov VV, Ren Q, Ferris P, Lindquist E, Shapiro H, Lucas SM, Grimwood J, Schmutz J, Cardol P, Cerutti H, Chanfreau G, Chen CL, Cognat V, Croft MT, Dent R, Dutcher S, Fernández E, Fukuzawa H, González-Ballester D, González-Halphen D, Hallmann A, Hanikenne M, Hippler M, Inwood W, Jabbari K, Kalanon M, Kuras R, Lefebvre PA, Lemaire SD, Lobanov AV, Lohr M, Manuell A, Meier I, Mets L, Mittag M, Mittelmeier T, Moroney JV, Moseley J, Napoli C, Nedelcu AM, Niyogi K, Novoselov SV, Paulsen IT, Pazour G, Purton S, Ral JP, Riaño-Pachón DM, Riekhof W, Rymarquis L, Schroda M, Stern D, Umen J, Willows R, Wilson N, Zimmer SL, Allmer J, Balk J, Bisova K, Chen CJ, Elias M, Gendler K, Hauser C, Lamb MR, Ledford H, Long JC, Minagawa J, Page MD, Pan J, Pootakham W, Roje S, Rose A, Stahlberg E, Terauchi AM, Yang P, Ball S, Bowler C, Dieckmann CL, Gladyshev VN, Green P, Jorgensen R, Mayfield S, Mueller-Roeber B, Rajamani S, Sayre RT, Brokstein P, Dubchak I, Goodstein D, Hornick L, Huang YW, Jhaveri J, Luo Y, Martínez D, Ngau WC, Otillar B, Poliakov A, Porter A, Szajkowski L, Werner G, Zhou K, Grigoriev IV, Rokhsar DS, Grossman AR. (2007). The Chlamydomonas genome reveals the evolution of key animal and plant functions. Science. 2007 Oct 12;318(5848):245–50.

Moseley JL, Page MD, Alder NP, Eriksson M, Quinn J, Soto F, Theg SM, Hippler M, Merchant S. (2002). Reciprocal expression of two candidate di-iron enzymes affecting photosystem I and light-harvesting complex accumulation. Plant Cell. 2002 Mar;14(3):673–88.

Prochnik SE1, Umen J, Nedelcu AM, Hallmann A, Miller SM, Nishii I, Ferris P, Kuo A, Mitros T, Fritz-Laylin LK, Hellsten U, Chapman J, Simakov O, Rensing SA, Terry A, Pangilinan J, Kapitonov V, Jurka J, Salamov A, Shapiro H, Schmutz J, Grimwood J, Lindquist E, Lucas S, Grigoriev IV, Schmitt R, Kirk D, Rokhsar DS. Genomic analysis of organismal complexity in the multicellular green alga Volvox carteri. Science. 2010 Jul 9;329(5988):223–6.

Strang, G. (2009). Introduction to Linear Algebra. Wellesley-Cambridge Press.

Zachariae W, Schwab M, Nasmyth K, Seufert W. Control of cyclin ubiquitination by CDK-regulated binding of Hct1 to the anaphase promoting complex. Science. 1998 Nov 27;282(5394):1721–4.

Zur H, Tuller T. (2013). New universal rules of eukaryotic translation initiation fidelity. PLoS Comput Biol. 2013;9(7):e1003136. doi: 10.1371/journal.pcbi.1003136. Epub 2013 Jul 11.

